# Spatial imprints of emergent cardiomyocyte states in the pressure-overloaded heart

**DOI:** 10.64898/2026.05.04.721738

**Authors:** Yuening Liu, Annabelle M. Coles, Jonah Castiglione, Vijay Venu Thiyagarajan, Kalen Clifton, Deevanshu Goyal, Jerry Wu, Arlo Sheridan, Ana Vujic, Kristen M. Harris, Uri Manor, Talmo D. Pereira, Jean Fan, Richard T. Lee, Pallav Kosuri

## Abstract

Resilience to cardiac stress is essential for health, yet the relationship between cardiomyocyte (CM) stress response and local microenvironment remains unclear. Here, we combined MERFISH spatial transcriptome profiling with *Cellouette*, an improved cell segmentation method, to determine CM-microenvironment relationships in a mouse model of ventricular pressure overload. We report the shape, transcription profile, spatial organization, and physical connectivity for >400,000 cells across stressed and healthy tissues. Under stress, CMs adopted a spectrum of emergent transcriptional states, with advanced states marked by a metabolic and pro-fibrotic shift. To discover CM-environment relationships, we performed a network analysis of physical cell connectivity combined with cell-type-specific profiling. We found that pro-fibrotic CM progression was tightly linked to distinct local microenvironments, and CM metabolic shifts could be inferred from transcriptional patterns in neighboring non-CM cells, revealing microenvironmental imprints of disease. We thus provide a resource for understanding the heterogeneity of outcome during cardiac pressure overload.

**Highlights:**

- Cellouette provides accurate segmentation for single-cell spatial transcriptomics in cardiac tissue.
- Pressure overload creates spatial gradients of cardiomyocyte pro-fibrotic states.
- Cardiomyocyte pro-fibrotic progression is linked to changes in local cell composition and gene expression.
- Transcriptional states of non-muscle cells predict metabolic state of adjacent cardiomyocytes.

## Introduction

Cardiomyocytes (CMs), the terminally differentiated muscle cells of the heart, are responsible for generating the contractile force required to maintain systemic blood circulation^1,2^. Unlike other cells in the heart, CMs have limited proliferative capacity, making their long-term integrity essential for preserving cardiac homeostasis and preventing functional decline^3–5^. Over a lifetime, CMs are subject to persistent mechanical stress, and this stress is exacerbated by common conditions such as hypertension, which affects nearly 50% of US adults^6^. Such persistent increase in pressure causes a gradual remodeling of the heart, resulting in hypertrophy and fibrosis, which eventually progresses to heart failure^7–9^. Strategies to better preserve cardiac health across the life span would therefore benefit from an understanding of the factors that regulate cardiomyocyte homeostasis under conditions of stress. However, the molecular and microenvironmental factors that determine CM health and resilience under pressure overload are not fully known.

Cardiac pressure overload triggers CM transcriptional and morphological remodeling, including metabolic reprogramming, hypertrophy, and impaired contractility ^8,10,11^. On the level of single cells, the responses of CMs to pathological stress are heterogeneous, as evidenced by the transcriptionally distinct CM states identified in animal models of heart disease and in human patients suffering from cardiac hypertrophy and heart failure^12–15^. However, most previous single-cell studies have relied on methods that require dissociation of cells from the tissue and thus come with a loss of the cells’ spatial context. The missing contextual information is significant because changes in a CM may be linked to its environment. For instance, distinct CM states are tightly tied to their local cellular microenvironment during development^16^ and during myocardial infarction^17^. In contrast, less is known about spatially and contextually distinct CM states in hearts under conditions of global stress, such as during pressure overload. Nonetheless, two lines of evidence indicate that pressure overload results in spatially and contextually distinct CM states. First, cardiac pressure overload leads to an increasingly high degree of transcriptional heterogeneity among individual cardiomyocytes^12,14,18^. Second, the cellular environments in a pressure overloaded heart progress at different paces, causing rapidly growing differences in the cellular and molecular environments surrounding individual CMs. Specifically, pressure overload initially causes perivascular fibrosis, which entails fibroblast expansion and activation, followed by excessive extracellular matrix (ECM) deposition in the areas immediately surrounding major blood vessels^19,20^. Later in the disease progression, fibrosis also spreads to the interstitial spaces between CMs in the myocardium. Thus, pressure overload leads to the emergence of heterogenous CM states as well as an emergence of heterogenous cellular and molecular microenvironments within the myocardium. We therefore hypothesized that under pressure overload, the state of a CM may be linked to its local microenvironment. Uncovering such connections would allow us to define environmental markers of CM dysfunction. By extension, these findings could aid in the discovery of novel drivers of CM health and resilience under pathological stress.

Here, we used a mouse model of cardiac pressure overload to determine the relationship between CM response and their cellular and molecular microenvironments. To do this, we first developed a workflow to enable measurement of the precise location, shape, and transcriptome profile of every cell in a cardiac tissue section (Figure 1A, Video S1&S2). Using this workflow, we profiled more than 400,000 cells in pressure overloaded hearts and in healthy controls. We then used our spatial transcriptomic map of cells to identify a disease-associated pro-fibrotic CM state and found that CMs in this state colocalized with conventional hallmarks of disease. To discover links between the state of a CM and its immediate environment, we then created a graph representation of physical cell connectivity. Using this connectivity graph and other approaches, we discovered cell-type-specific microenvironmental changes that were associated with distinct responses in the local CMs. We found that though different hearts displayed different spatial patterns of disease progression, the underlying spectrum of transcriptional states sampled by the CMs was remarkably similar across hearts. Our observations led us to hypothesize that CM state might be consistently linked to the transcriptional states of local non-CM cells. To test this hypothesis, we trained a neural net to predict CM progression based on transcriptional profiles of neighboring non-CM cells. Remarkably, we found that the transcriptional states of non-CM cells could be used to reconstruct the spatial patterns of CM progression in early-stage heart disease. Our results thus show that during cardiac pressure overload, CM state is tightly coupled to distinct and cell-type specific gene expression patterns in neighboring non-CM cells.

**Figure 1.**
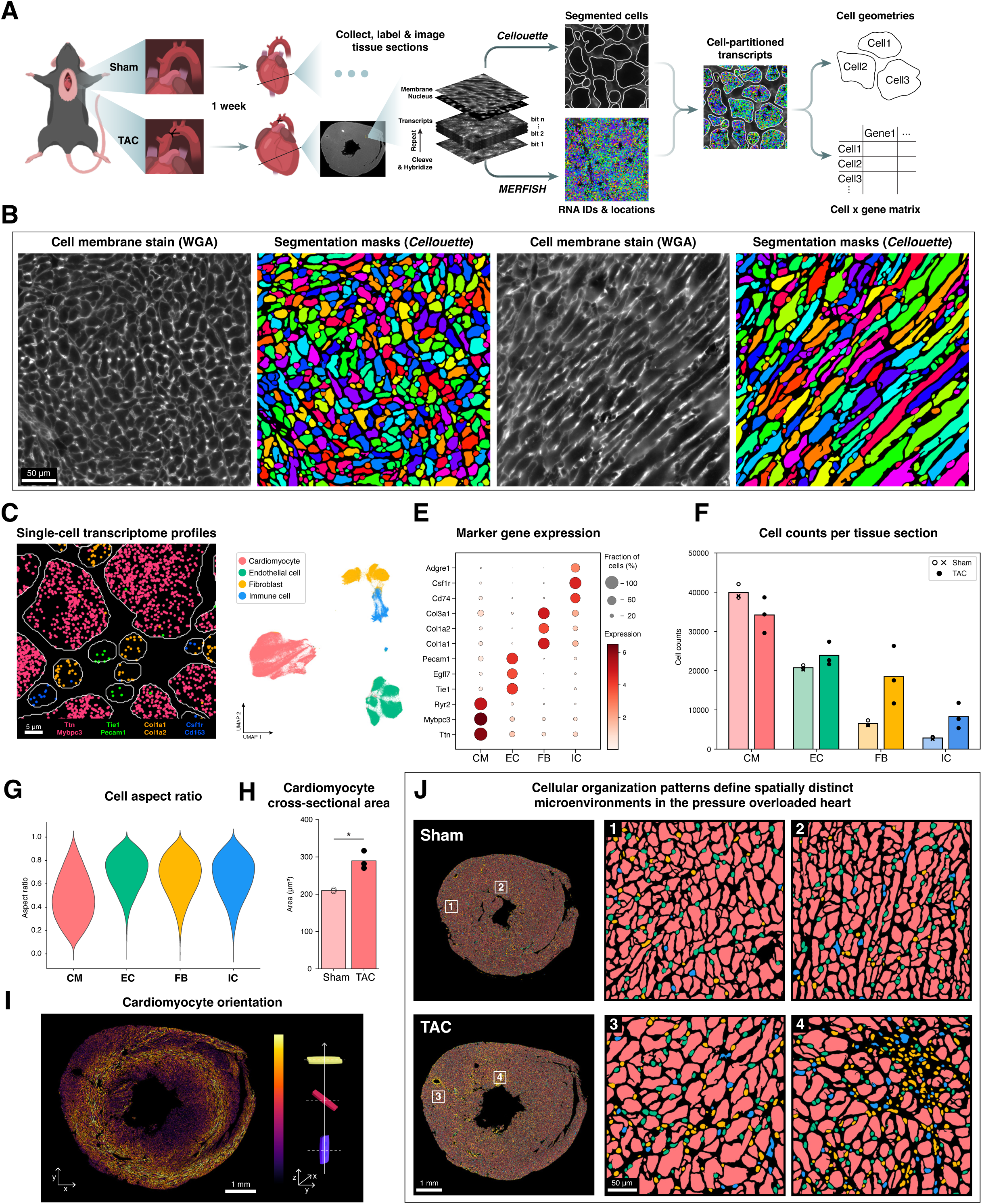
Cellouette enables single-cell spatial profiling in healthy and pressure overloaded mouse hearts. (A) Experimental workflow. Mice were either subjected to transverse aortic constriction (TAC; *N*=3) or sham surgery (sham; *N*=3). Hearts were collected one-week post-surgery and immediately frozen. Mid-ventricular transverse sections 10 µm thick were fixed, labeled for MERFISH imaging, embedded in a hydrogel, and chemically cleared. In each section, hundreds of fields of views were imaged and computationally stitched together. Transcripts were detected in sequential rounds of imaging and then decoded to reveal their identity and spatial location. Cell boundaries were stained using wheat germ agglutinin (WGA), nuclei using DAPI, and the images were segmented using Cellouette. Based on segmentation-defined cell boundaries, transcripts were then partitioned to individual cells, yielding single-cell locations, geometries, and transcriptional profiles. (B) Cellouette provides high-performance segmentation of diverse cell shapes in crowded tissue. Representative fields of view showing WGA-stained cell boundaries and Cellouette cell segmentation in myocardial locations with cross-sectional (*left panels*) and longitudinal (*right panels*) cardiomyocyte orientations. (C) Validation of cell segmentation and transcript partitioning accuracy. Representative field of view showing spatial segregation of cell-type-specific transcripts, indicating a low level of cross-contamination of transcripts between neighboring cells. Marker genes were chosen for major cell types: Ttn/Mybpc3 (cardiomyocytes), Tie1/Pecam1 (endothelial cells), Col1a1/Col1a2 (fibroblasts), Csf1r/Cd163 (macrophages / immune cells). (D) Classification of major cell types. Combined clustering of the sham and TAC cells (*n*=425,027) displayed in a UMAP. Major cell types are shown, including cardiomyocyte (CM), endothelial cell (EC), fibroblast (FB), and immune cell (IC). (E) Validation of cell type classification. Major cell types show mutually exclusive expression of canonical marker genes. (F) Cell population analysis of the major cardiac cell types in sections from sham (*N*=3) and TAC (*N*=3) hearts. Data points indicated with a cross indicate values from a partial sample (see methods). (G) Distribution of cell aspect ratios (short axis / long axis) for the four major cardiac cell types. (H) Average area of cross-sectioned CMs in sham (*N*=3) and TAC (*N*=3) heart sections. Mean ± s.d., \**p* < 0.05 (unpaired two-tailed t-test). (I) Spatial distribution of CM orientation in a TAC heart section, with color indicating the magnitude of the out-of-plane orientation component. (J) Emergent cellular microenvironments in the pressure overloaded heart. In the sham heart (*top row*), non-CM cells are seen dispersed and at similar densities throughout the tissue section. Two representative areas are shown in magnified view. In the TAC heart (*bottom row*), some areas display dispersed non-CM cells reminiscent of the organization in healthy hearts (*left inset*), while other areas (*right inset*) reveal local clusters of FBs and ICs, indicating emergence of distinct cellular microenvironments in the 1-week TAC heart.

## Results

### Cellouette and MERFISH enables single-cell spatial profiling of the heart during pressure overload

Pressure overload was induced using transverse aortic constriction (TAC)^21^, a widely used mouse model that generates hemodynamic stress and promotes cardiac remodeling, reproducing key features of hypertension-induced heart failure in humans^22^. To validate the physiological response to TAC, echocardiographic assessments were performed at baseline (prior to surgery) and one-week post-surgery (Figure S1A-B). No significant differences were observed between animal groups at baseline. However, at one-week post-surgery, TAC mice (*n*=3) exhibited reduced fractional shortening (FS) and increased interventricular septal thickness in diastole (IVSd), posterior wall thickness in diastole (IVPWd), and left ventricular mass in diastole (LVMd), compared to sham-operated control mice (*n*=3), indicating systolic dysfunction and ventricular hypertrophy. These observations confirmed effective induction of pressure overload resulting in ventricular remodeling and the onset of cardiac disease.

To investigate the impact of pressure overload on gene expression in the ventricular myocardium, we collected hearts from TAC and sham mice at one-week post-surgery, immediately froze the dissected hearts, and sectioned them along the short axis at the mid-ventricular level. One section from each heart was then processed for multiplexed error-robust fluorescence *in situ* hybridization (MERFISH)^23,24^ (Figure 1A). To identify key cell types and investigate processes relevant to the pathophysiology of pressure overload, we designed, synthesized, and labeled the cardiac tissue samples with a custom probe set targeting 128 genes selected to contain markers for major cell types and disease pathways (Table S1). MERFISH imaging was performed on a custom-built microscope setup with automated fluidics, stage control, and autofocus, to perform sequential rounds of hybridization and imaging. Through image analysis, we then detected, localized, and decoded tens of millions of individual RNA molecules within each tissue section. Our MERFISH data showed reproducibility between biological replicates, and total transcript counts showed high correlation with bulk RNA sequencing data^25^, validating our *in situ* measurements of gene expression (Figure S1C).

Accurate profiling of single cells *in situ* requires accurate cell segmentation, however cardiac tissue is densely packed, structurally complex, and composed of cells with heterogenous shapes, making automated single-cell segmentation particularly challenging. To aid in image-based segmentation, we stained nuclei using DAPI and cell boundaries using wheat germ agglutinin (WGA), which binds to glycoproteins on the cell surface, and then imaged these labels during MERFISH data acquisition. The WGA-labeled images revealed clear cell boundaries in every section (Figure 1B). Manual annotation of these boundaries served as a ground-truth reference for automated cell segmentation. We first evaluated several commonly used cell segmentation approaches (Figure S1D-G). DAPI-seeded watershed segmentation did not yield useful results as the method relies on the assumption that nuclei are located at the centers of cells. Rather, we found that for many cells, the nucleus was not captured within the section or was off-center, and large multinucleated cardiomyocytes often became split into multiple segments, resulting in inaccurate segmentation. We also evaluated Cellpose SAM and a Cellpose 2.0 model^26^ custom-trained on our manually annotated ground-truth reference. We found that the Cellpose models performed well on small non-CM cells and on some cross-sectioned cardiomyocytes but failed to segment longitudinally oriented cardiomyocytes (Figure S1D). The Cellpose algorithm computes pixel-wise vector fields where each pixel’s vector points towards the center of the cell in which the pixel is located. Cellpose then generates segmentations by grouping pixels whose vectors converge to a common point, indicating that these pixels belong to the same cell. However, for large, elongated, and non-convex cells, there is often no central point of convergence across the entire cell geometry, resulting in fragmented, missing, or inaccurate segmentations. To address these limitations, we developed Cellouette, a segmentation pipeline optimized to accurately segment cells of a wider range of geometries such as large and irregular cardiomyocytes, in densely packed tissue. Cellouette combines the use of a nuclear Cellpose model to segment small cells (with minimal cytoplasm) based on their DAPI signal, with a new 3D segmentation model that segments larger cells based on their WGA signal. Our new model predicts direct neighbor voxel-wise affinities and improve these affinities through prediction of local shape descriptors (LSDs), which provide additional contextual information by describing the local shape of the cell boundary^27^. Cellouette enabled superior segmentation accuracy in tissue containing both cross-sectional and longitudinally oriented CMs, as well as smaller cells within crowded myocardial environments (Figure 1B, Figure S1D-G). Using the Cellouette segmentation boundaries, we assigned our spatially resolved transcripts to the segmented cells, enabling measurements of single-cell shapes, positions, and transcription profiles across the tissue sections. In total, our analysis yielded profiles of more than 400,000 cells across the sections from sham and TAC hearts.

A particularly problematic consequence of segmentation errors is the misassignment, or “leakage”, of transcripts between neighboring cells. Transcript leakage complicates the discovery of cell-neighborhood relationships because it artificially makes cells appear more transcriptionally similar to their neighbors. To evaluate our accuracy of transcript assignment, we therefore measured the co-expression of mutually exclusive marker genes for distinct cardiac cell types across all segmented cells (Figure 1C, Figure S1H). Minimal co-expression was observed between marker genes for different cell types, indicating minimal transcript leakage and confirming the accuracy of both cell segmentation and transcript assignment.

To identify cells according to the major cardiac cell types, and to measure population-level changes in cell numbers upon pressure overload, we computationally integrated our single-cell MERFISH expression data from all tissue sections across conditions and replicates, classified the cells using batch-corrected unsupervised clustering, and visualized these clusters in a UMAP (Figure 1D, Table S2). Four major clusters were identified based on expression of canonical marker genes (Figure 1E), corresponding to cardiomyocytes (CM), endothelial cells (EC), fibroblasts (FB), and immune cells (IC). Pressure overload led to differences in both cell abundance and overall cellular composition (Figure 1F, Figure S1I). Specifically, we observed an increase in the number of FBs and ICs in TAC hearts, consistent with the fibrotic and inflammatory response in early-stage pressure overload^28–30^.

Increased ventricular load causes concentric hypertrophy due to the growth in volume of individual cardiomyocytes^8^. To quantify cardiomyocyte hypertrophy and control for CM orientation bias within a tissue section, we considered cell aspect ratios (Figure 1G) to distinguish between cross-sectional and longitudinal cardiomyocytes. In this way, we could accurately compare cardiomyocyte size between conditions. We found that CMs had ∼1.4x larger cross-sectional areas in TAC *vs*. sham (Figure 1H), indicating substantial hypertrophic growth in response to pressure overload, similar to previous reports^31^.

The muscle fibers in the ventricular myocardium are oriented in a variety of directions. While fiber stress is largely even across the myocardium in a healthy heart, pressure overload may result in different levels of stress in CMs with different orientation angles^32^. However, investigations into the link between CM orientation and stress response have been hindered by the challenge of identifying local myofiber architecture directly in cardiac sections. To overcome this challenge, we developed an orientation field approach based on CM segmentation geometries to directly map local myofiber 3D orientation within our sections (see Methods). Our resulting single-cell spatial maps reveal the patterns of myofiber orientation within the ventricular myocardium (Figure 1I).

Pressure overload led to changes in the relative abundance of different major cell types (Figure 1F), which by necessity resulted in a change in the local microenvironments of individual CMs. To examine if these changes were spatially localized, we examined the spatial distributions of cell types within the sham and TAC hearts. In the sham hearts, we found that the major cell types displayed an even pattern without much variation across the myocardium (Figure 1J, *areas 1 and 2*). The TAC hearts also contained regions with evenly distributed cells (Figure 1J, *area 3*), though notably with a higher density of FBs and ICs, consistent with our cell population-level measurements. However, certain regions within the TAC hearts displayed a breakdown of this order, including areas with high FB density and areas with an absence of CMs where instead clusters of FBs and ICs were found, likely indicating sites of replacement fibrosis (Figure 1J, *area 4*). Thus, in contrast to the evenly distributed cellular patterns observed in the sham hearts, the TAC hearts revealed a heterogeneous and clustered distribution of major cell types across different myocardial regions.

Taken together, our results provide spatial single-cell maps of healthy and pressure overloaded ventricular myocardia, and reveal the emergence of localized cellular heterogeneity upon pressure overload. We thereby demonstrate that by combining Cellouette with MERFISH, we can uncover both global and local changes in the composition, orientation, structure, and organization of cells in the mammalian heart.

### Emergence of stress-induced cardiomyocyte states in the pressure-overloaded heart

In a healthy heart, CMs in the left ventricular wall have no major functional heterogeneity. However, CMs can be affected by changes in their cellular environment^33^, and pressure overload causes the emergence of new and distinct cellular environments, as evidenced by our single-cell maps (Figure 1J). We therefore hypothesized that these stress-induced microenvironments may be associated with altered CM states during pressure overload.

To identify stress-induced CM states, we analyzed transcript counts within CM boundaries (Figure 2A). Canonical CM markers such as the structural sarcomeric gene titin (Ttn) were consistently expressed in both sham and TAC hearts. By contrast, CMs from TAC hearts showed an emergent and heterogeneous expression of genes linked to hypertrophy (Acta1) and fibrosis (Col1a1, Col1a2, Col3a1). Collagen expression is typically associated with FBs, although some previous evidence has pointed to CMs also expressing collagen in pressure overloaded hearts^34,35^. False positive findings are common in single-cell studies as transcriptome measurements may erroneously include transcripts from neighboring cells. This risk is especially high in spatial single-cell studies, where correct assignment of transcripts to cells depends on accurate cell segmentation. Since collagen-expressing FBs are abundantly present in pressure overloaded hearts, even minor errors in cell segmentation could cause assignment of FB-derived transcripts to adjacent CMs, which in turn would lead to the false discovery of collagen expression in CMs. Therefore, to verify that our CM-assigned collagen transcripts were indeed found within CMs, we inspected the spatial distribution of transcript localizations relative to cell boundaries both from our segmentation data and the corresponding fluorescence images. As seen in Figure 2A, collagen transcripts were clearly found in levels well above background throughout the CM cell geometries in the TAC condition, thus confirming that pressure overload leads to transcription of collagen genes in CMs.

**Figure 2.**
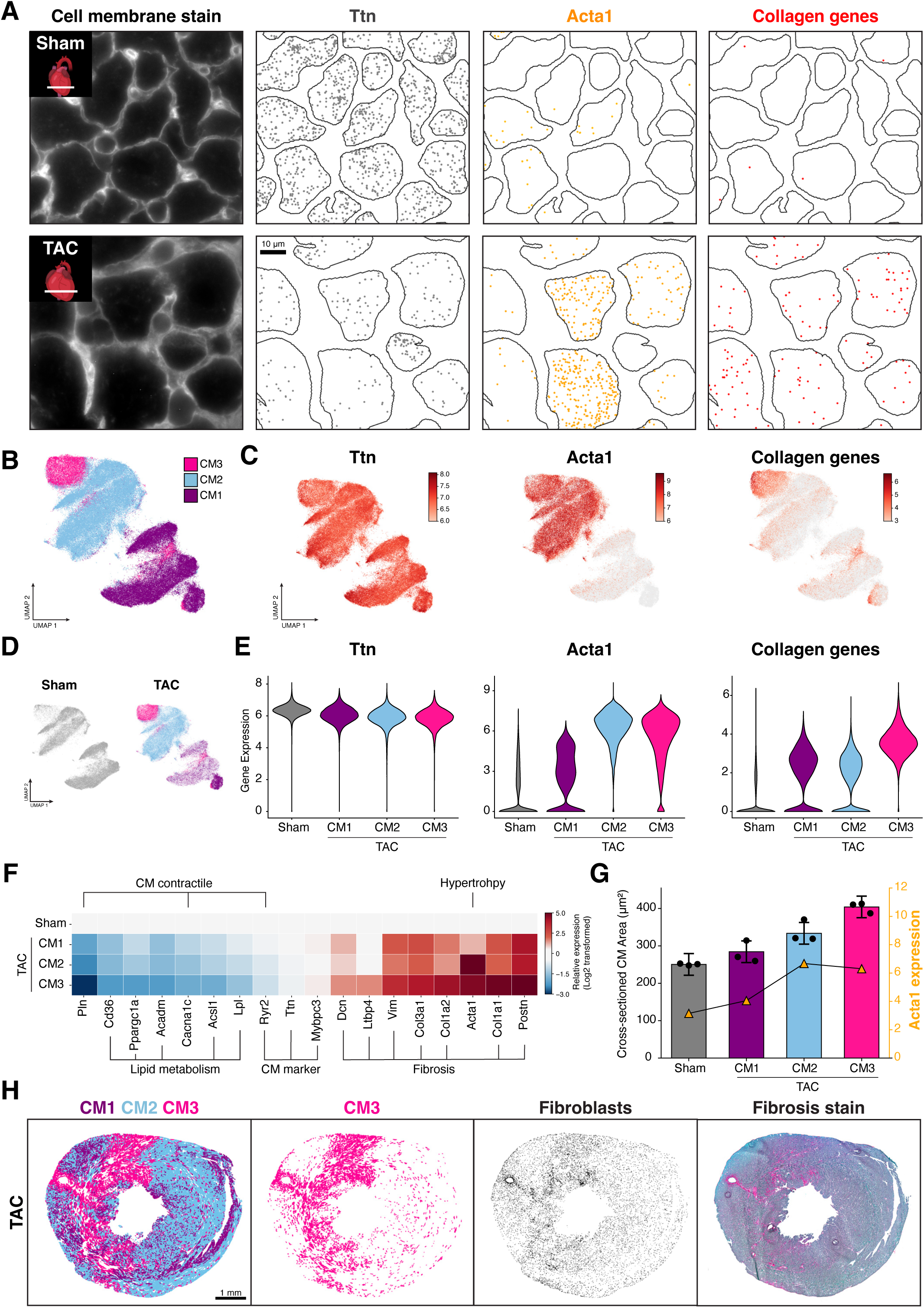
Pro-fibrotic transformation of cardiomyocytes in the pressure overloaded heart. (A) CM transcript densities in the ventricular myocardium from sham (*top row*) and TAC (*bottom row*) hearts. Leftmost image shows fluorescent cell membrane stain (WGA). Following images show corresponding segmented CM cell boundaries and their respective transcripts from marker genes of CM identity (Ttn), CM hypertrophy (Acta1), and fibrosis (Collagen genes: Col1a1, Col1a2, Col3a1). (B) UMAP showing semi-supervised clustering of CMs (*n*=147,761) from sham and TAC hearts. (C) Expression maps for the genes shown in (A) across all CMs. Color indicates transcript density. (D) The UMAP of CMs from sham and TAC hearts, respectively. Sham CMs (gray) referred to together as a single group of healthy cells, while TAC CMs referred to according to the clusters CM1 (purple), CM2 (blue), and CM3 (magenta). (E) Gene expression distributions in the sham CMs and in the three CM clusters in TAC. (F) Gene expression in the three TAC CM clusters relative to sham. (G) Geometric and transcriptional measures of CM hypertrophy. Bars and black circles show the average area of cross-sectioned CMs (*left axis*). Each point represents the average from a single heart. Line and yellow triangles show average Acta1 expression (*right axis*) in the corresponding CMs. (H) Pro-fibrotic CMs (CM3) colocalize with FBs and with sites of tissue fibrosis. *From left*: Spatial distribution of CM states in a TAC heart; location of pro-fibrotic CMs (CM3); location of FBs; fibrosis histology stain (Sirius red) of an adjacent tissue section. Gene expression values in (C), (E), and (G) are normalized to cell volume and log-transformed.

Pressure overload caused a diversification of transcriptional states among cardiomyocytes. To define these states, we performed semi-supervised clustering on the gene expression profiles from the 147,761 CMs in sham and TAC conditions combined (Figure 2B-D). Our clustering analysis revealed three major, distinct CM subpopulations that we refer to as CM1, CM2, and CM3. Of these three populations, CM1 was mainly represented in sham hearts, CM2 was mainly found in TAC hearts, and CM3 was almost exclusively found in TAC hearts (Figure 2D). All three subpopulations expressed the CM marker gene Ttn (Figure 2C). Both CM2 and CM3 expressed high levels of Acta1, potentially indicating CM hypertrophy. CM3 additionally expressed collagen genes (Col1a1, Col1a2, Col3a1) and periostin (Postn), indicating that in the pressure overloaded hearts, a fraction of the hypertrophic CMs had adopted a pro-fibrotic state (Figure 2C-E, Figure S2A).

Comparison of gene expression profiles between sham CMs and the three TAC CM populations revealed differences in gene expression across several functional domains (Figure 2F; we hereafter grouped all sham CMs together as a separate healthy reference). Compared with sham CMs, TAC CMs had lower expression of genes involved in fatty acid (FA) metabolism (eg. Cd36, Acsl1, Acadm, Lpl) consistent with the shift from oxidative phosphorylation to glycolysis seen in CMs under excessive stress^36^. We also found decreased expression of genes related to calcium handling (eg. Cacna1c, Pln, Ryr2), potentially indicating impaired excitation-contraction coupling. Meanwhile, fibrosis-associated genes were expressed in higher amounts (eg. Col1a1, Col1a2, Col3a1, Postn), showing a pro-fibrotic shift in CM state that included expression of genes typically associated with activated fibroblasts. This pro-fibrotic shift was seen in all TAC CM populations but was most pronounced in CM3 (Figure 2F). In fact, for nearly all the genes that showed differential expression between sham and TAC, CM3 had the expression value furthest from the healthy sham cells, suggesting that CM3 constituted the most progressed population of cells. Meanwhile, expression of adult sarcomeric genes (eg. Ttn, Mybpc3) remained mostly unchanged across conditions and populations, indicating an absence of systematic measurement errors or batch effects. Overall, we found that CM transcription profiles were more varied in TAC *vs.* sham hearts (Figure S2B), indicating that pressure overload leads to the emergence of new and heterogenous CM states.

Of the three CM populations in the TAC hearts, CM3 was transcriptionally the most different from sham. However, the hypertrophic marker gene Acta1 was a notable exception, as its transcript density appeared lower in CM3 than in CM2 (Figure 2G, *right axis*). To test if the CM3 cells were indeed less hypertrophic than the CM2 cells, we compared the cross-sectional area of cardiomyocytes from each group, using the cell geometry analysis introduced in Figure 1H. Surprisingly, and in contrast to our measurements of Acta1 expression, we found a progressive increase in cell size across CMs from sham, CM1, CM2, and CM3, respectively, with CM3 displaying the largest cross-sectional CM areas (Figure 2G, *left axis*). These results strengthen our conclusion that CM3 represents the most progressed population of CMs. Furthermore, these findings show that Acta1 expression alone cannot be reliably used to determine the extent of CM hypertrophy.

Our finding that CM3 constituted the cells most severely affected by pressure overload led us to ask, where in the ventricular myocardium these cells were located. If these cells were spatially enriched in certain regions of the ventricle, then that could indicate an association with distinct disease-associated microenvironments. Indeed, we found that CM3 cells were mainly found in spatially contiguous regions (Figure 2H). Since the CM3 cells displayed pro-fibrotic expression profiles, we wondered if the CM3 regions might be defined by broader fibrotic features in the tissue. To answer this question, we performed two comparisons. First, to see if the CM3 cells colocalized with sites of increased fibroblast abundance, we created a map of all the fibroblasts in the section. We found that the pattern of increased fibroblast density indeed resembled the pattern of CM3 cells (Figure 2H). Second, we collected a tissue section adjacent to the section used for MERFISH, and stained this adjacent section with Sirius Red to identify sites of local tissue fibrosis. After alignment of the histology-stained section with the MERFISH section and computationally extracting the fibrotic areas (Figure S2C), we found that in the TAC heart, CM3 cells colocalized with the most severely fibrotic areas of the tissue (Figure 2H). This was not observed in the sham heart (Figure S2D). We thus conclude that the most progressed CMs are located in regions of increased fibroblast abundance and excessive ECM deposition.

In summary, we found that CMs in the pressure overloaded hearts shifted into states of varying progression. CMs in the most progressed of these states, CM3, displayed the lowest expression of FA metabolism and calcium handling genes, the highest degree of hypertrophy, and a shift to a pro-fibrotic phenotype that included expression of collagen and periostin. These pro-fibrotic CMs were found in the most fibrotic areas of the tissue, and were thus associated with large-scale cellular, chemical, and mechanical changes in their local environment.

### Spatial patterns of cardiomyocyte-specific gene expression during pressure overload

Our previous analysis relied on the clustering of CMs into discrete states based on expression of many genes. However, such discrete classification can sometimes obscure differentiation of cells along multiple gene expression axes that are not aligned. To test whether the spatial patterns of pro-fibrotic CMs represented patterns of disease progression across other functional axes, we therefore performed on a TAC heart section a spatially variable gene (SVG) analysis that did not rely on clustering CMs into discrete states (Figure 3, Figure S3). Our SVG analysis led to the identification of genes that had a high spatial autocorrelation and a statistically significant variation in CMs across the tissue (see Figure 3A for two examples of such genes). We then measured the cross-correlation of these SVGs and discovered groups of genes that had similar spatial patterns of expression in CMs within a heart section. Strikingly, we found that the top two groups of similar-patterned genes had a strong within-group correlation and a strong anticorrelation between groups (Figure 3B, Figure S3A). The first group (*pattern A* in Figure 3B) included genes involved in fibrosis, and the second group (*pattern B* in Figure 3B) included genes involved in FA metabolism. These results support our previous finding that pro-fibrotic CMs had lower expression of FA metabolism genes (Figure 2F) and show that these two groups of genes (pro-fibrotic and FA metabolism) share a single axis of CM differentiation.

**Figure 3.**
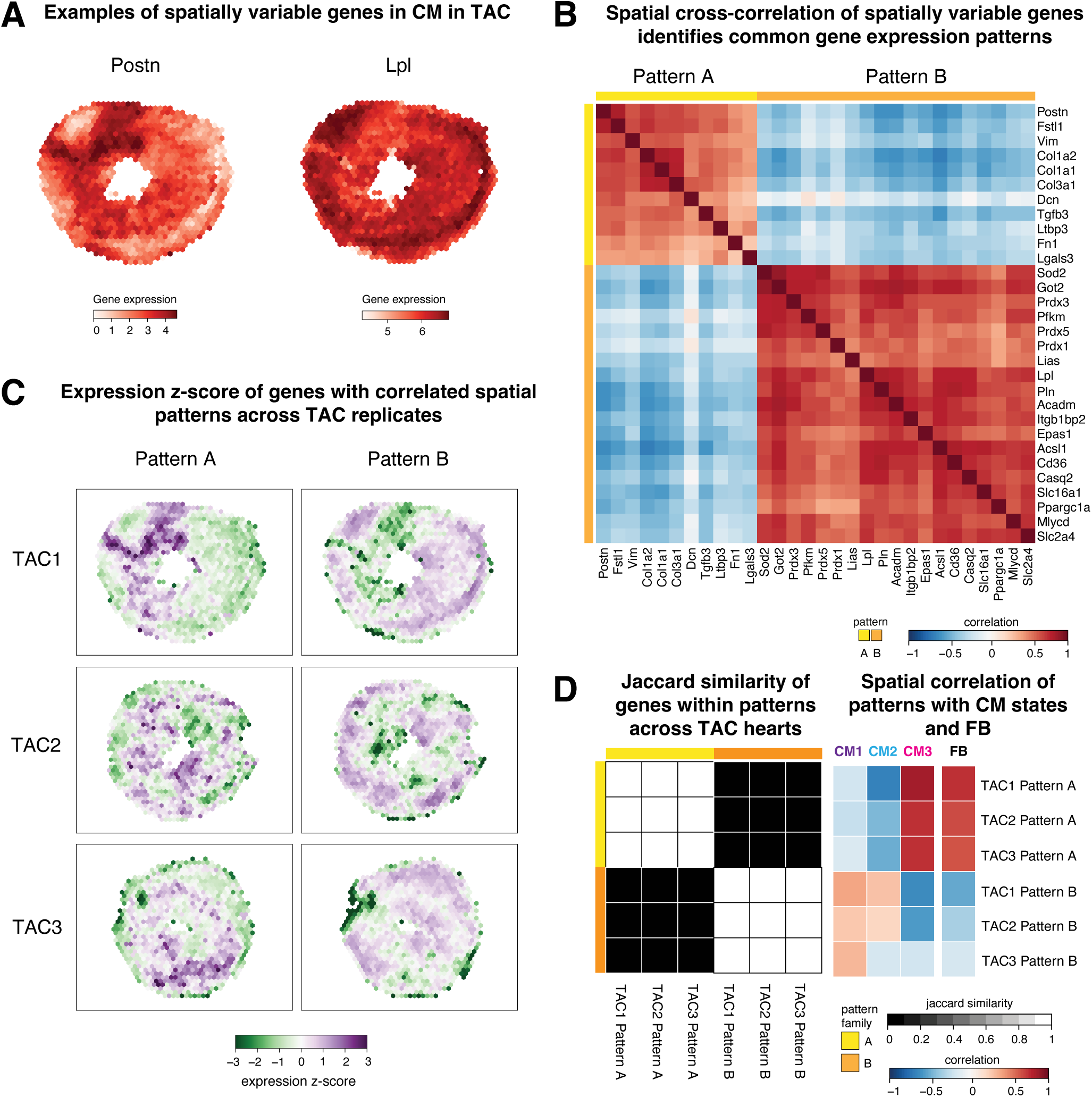
Spatially variable gene (SVG) patterns reveal conserved CM microenvironment programs across TAC hearts. (A) Examples of gene expression patterns in CMs for two of the 87 genes identified as SVG in a TAC heart. Gene expression of CMs normalized to cell volume and then rasterized into 171 μm hexbins. Color indicates mean volume-normalized expression of CMs within each hexbin, log-transformed. (B) Correlation matrix of the hexbin gene expression for 30 SVGs in a TAC heart which cluster into two spatial patterns groups (A and B). (C) Expression of genes within each spatial pattern group (*left column*: A, *right column*: B) for each TAC heart (*top row*: TAC1, *center row*: TAC2, *bottom row*: TAC3) averaged and standardized using z-score normalization. (D) Genes within spatial patterns groups for all TAC hearts compared via Jaccard similarity (*left*). For each pattern, correlation of gene expression z-score with the spatial distributions of CM states (CM1–CM3) and with FB in the TAC heart sections to which the spatial pattern belongs (*right*).

To determine if these spatial gene groupings were consistent across biological samples, we measured the cross-correlation of the expression of the genes in patterns A and B in two other TAC hearts. We identified that these genes grouped into two distinct spatial patterns of expression in these hearts as well (Figure S3B). Although the spatial patterns of the gene groups were different in the different hearts (Figure 3C), the co-expressed genes within each group were identical across samples (Figure 3D, *left panel*). These results indicate that while CMs in different hearts may have different spatial patterns of progression, the transcriptional changes associated with CM progression are conserved across hearts.

We wondered how these spatial expression patterns were related to our previously identified pro-fibrotic CMs (CM3). We therefore measured the correlation of the SVG patterns with the spatial patterns of CM states and FB density (Figure 3D, *right panel*). These measurements showed that in all three hearts CM3 and FB density correlated strongly with Pattern A. Since Pattern A was marked by high expression of pro-fibrotic genes and low expression of FA metabolism genes, we conclude that the pro-fibrotic state (CM3) classification accurately captures the most progressed cells in the CM population, along both fibrotic and metabolic axes of differentiation. Taken together, our analyses in Figures 2 and 3 indicate that pressure overload induces regionally coordinated gene-regulatory programs in CMs, linking local microenvironmental cues to activation of pro-fibrotic genes and concurrent suppression of genes required for FA metabolism.

### Pro-fibrotic cardiomyocytes are associated with distinct cellular and molecular microenvironments

Interactions between CMs and other cell types are essential for cardiac homeostasis, and dysregulated interactions can contribute to the development and progression of heart disease^37–39^. Here, we took advantage of the spatial heterogeneity of CM states in the pressure-overloaded hearts to identify microenvironmental factors that were linked to pro-fibrotic CM progression.

Direct cell-cell connections mediate juxtacrine and electrical signaling between cells. To investigate differences in direct cellular connections of CMs, we constructed connectivity graphs based on physical proximity between neighboring cells, as determined from our segmented cell boundaries (Figure S4A). Although we could only detect cell connections within the plane of sectioning, we reasoned that due to the diversity of CM orientations sampled within our section (Figure 1I), any detected patterns would likely be representative of the broader tissue context. In the sham heart, our connectivity graph revealed a nearly completely interconnected network of CMs interspersed with sparsely but evenly distributed cells from other populations (Figure 4A, Figure S4A-B). Notably, the local network connectivity appeared qualitatively similar across the entire tissue section from the sham heart. Meanwhile, the TAC heart revealed patchy and less interconnected networks that varied substantially in composition and connectivity across the tissue section, indicating both an overall breakdown in cell-cell connectivity and an increase in heterogeneity (Figure 4A, Figure S4A-B). To quantify these changes, we calculated the average number of cell-type-specific connections per CM. We found that compared to the sham heart, CMs in the TAC heart had increased interactions with non-CM cells, particularly FBs and ICs (Figure 4B), consistent with the global increase in these cell populations (Figure 1F). When comparing the different CM states in the TAC heart, we found a trend from CM1 to CM3 showing an increasing severity as their connectivity to other cells increasingly differed from that of healthy CMs. Specifically, the pro-fibrotic CM3 cells had the highest number of connections with ICs and FBs, with both of these connections increasing 3-fold relative to sham (Figure 4B). These results corroborate our finding that the most severely progressed CMs are found in areas with the most progressed tissue fibrosis (Figure 2H) and reveal a link between the internal disease state of a CM and the concentration of ICs and FBs in its local cellular neighborhood.

**Figure 4.**
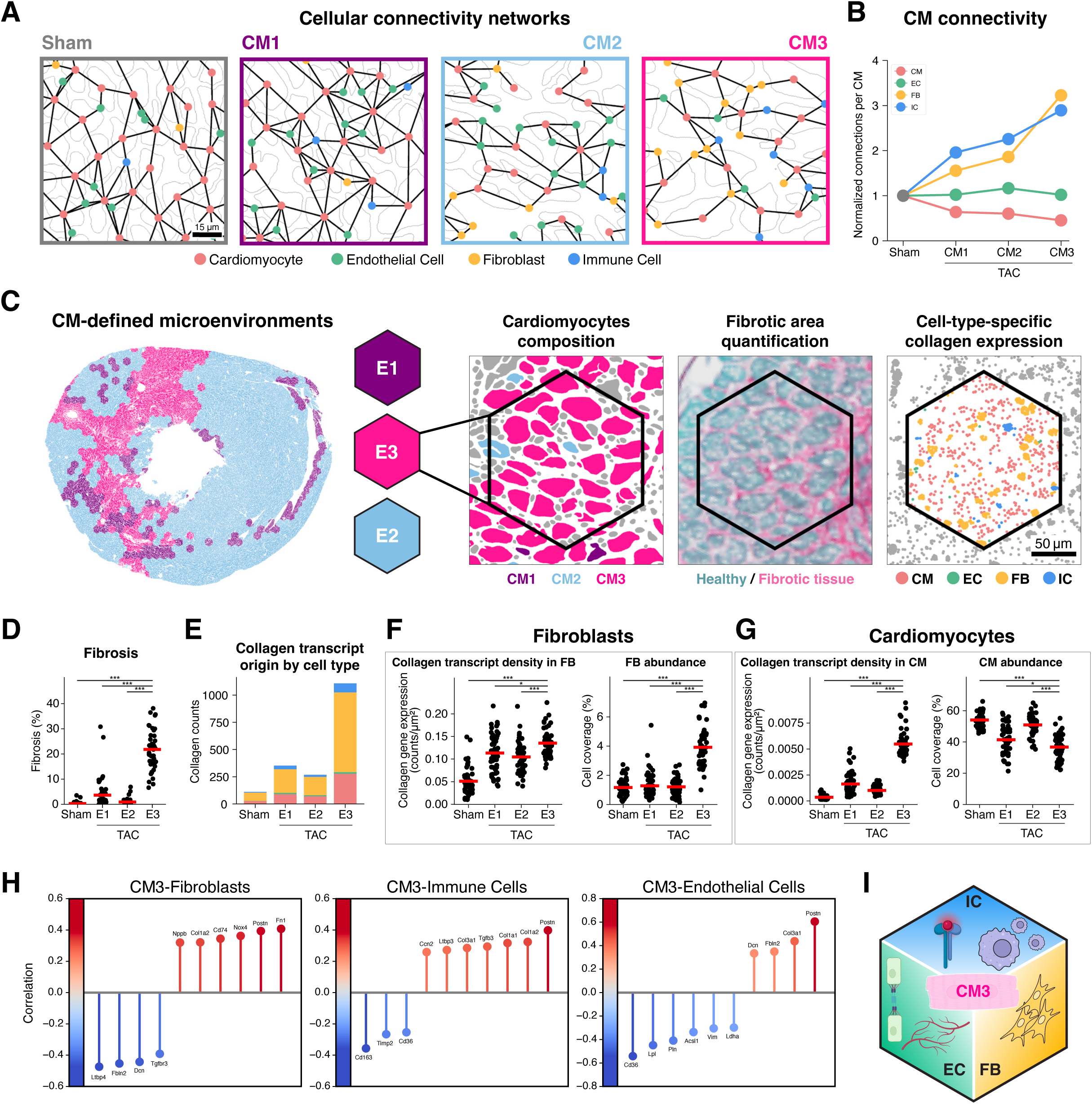
Pro-fibrotic cardiomyocytes (CM3) are associated with distinct microenvironments. (A) Spatial graphs showing observed physical connections between cells. Points indicate cell centroids and are colored according to major cell type. CM cell boundaries are shown in gray. Cells were considered connected when the shortest distance between the boundaries of two segmented cells was less than 3 μm. (B) CM connections observed with other CMs (*red*) and with non-CM cells (*green, yellow, blue*) for sham and for the three cardiomyocyte states in TAC. Connections per CM shown normalized to sham values. (C) CM microenvironment analysis pipeline. Single-cell maps and histology staining (Sirius red) images from a TAC section were computationally aligned and rasterized into a set of hexagonal bins. Each hexbin represents a local microenvironment and is classified as E1, E2, or E3 corresponding to its most enriched cardiomyocyte state (CM1, CM2, or CM3, respectively). Microenvironment measurements were made including composition of CM states, percentage of area covered with fibrosis, and cell-type-specific expression of collagen genes. (D) CM3-enriched microenvironments have the most area covered by fibrosis. Points represent individual microenvironments (*n*=50 per group); red lines show mean fibrosis percentage. *** p < 0.001 (two-sided Mann-Whitney U test). (E) Average number of collagen gene transcripts per hexbin in the four major cell types in the four microenvironments. (F) FB-specific collagen gene expression (*left*), and average FB area coverage (*right*) in each microenvironment (*n*=50 per group). Red lines show mean – values. * p < 0.05, *** p < 0.001 (two-sided Mann-Whitney U test). (G) CM-specific collagen gene expression (*left*), and average CM area coverage (*right*) in each microenvironment (*n*=50 per group). Red lines show mean values. * p < 0.05, *** p < 0.001 (two-sided Mann-Whitney U test). (H) Microenvironmental markers of pro-fibrotic CMs. Correlation between CM3 enrichment in a hexbin and the expression of individual genes in FBs (*left*), ICs (*middle*), and ECs (*right*) across each microenvironment. The 10 genes with the greatest absolute correlation coefficients are shown for each cell type. (I) Summary schematic illustrating the molecularly distinct microenvironment, or “imprints”, surrounding CM3 cardiomyocytes. ECs show signs of impaired lipid uptake, decreased angiogenic potential, and reduced cell-cell adhesion. FBs show signs of activation, TGF-β–driven proliferation, and differentiation towards myofibroblasts. ICs express more pro-inflammatory cytokine receptors and exhibit increased interleukin-mediated signaling.

Next, we asked if CMs in different states of pro-fibrotic progression were spatially associated with local tissue fibrosis and changes in gene expression in non-CM cells. To answer these questions, we computationally divided the tissue sections into hexagonal bins (hexbins) and assigned to each hexbin a label indicating its most common CM state (Figure 4C, Figure S4C). Hexbins in the TAC heart that contained mostly CM1 cells were thus labeled as E1, etc. For a healthy reference, we used hexbins from a sham heart. For each hexbin we could then relate CM progression to local fibrosis and cell-type-specific changes in gene expression (Figure 4C, Figure S4C).

Our hexbin analysis revealed that in the TAC heart, overt tissue fibrosis was highly localized to the E3 (i.e. CM3-enriched) microenvironments, while the E2 and E1 microenvironments were not significantly different than sham (Figure 4D). The concentration of fibrotic tissue in the E3 regions was reflected in the distribution of collagen transcripts that were also enriched in E3 (Figure 4E). To investigate the cell-type origin of these transcripts, we measured the contribution of each cell type to the overall expression of collagen. Unsurprisingly, we found that FBs were the dominant source of collagen transcripts across all microenvironments. However, to our surprise we found that CMs contributed a sizable fraction of collagen transcripts, amounting to nearly a third of the total partitioned transcripts in the E3 regions (Figure 4E).

Local increases in collagen expression can be due to upregulation of collagen within cells; however, it can also be due to proliferation of collagen-expressing cells. Measuring the relative contribution of these two mechanisms in a local microenvironment has historically been challenging. Here, we show that our approach allows us to cleanly disentangle the contribution from each mechanism. To determine the relative contributions of upregulation *vs.* proliferation to the expression of collagen, we measured the local density of collagen transcripts within FBs and CMs and also measured the local abundance of each cell type (Figure 4F-G). For FBs in low-fibrosis areas of the TAC heart (E1&E2), collagen transcript density was ∼2x increased relative to sham, while the FB abundance was unchanged. Thus, for FBs in low-fibrosis regions of TAC, the overall increase in collagen expression was mainly due to upregulation rather than proliferation. In the high-fibrosis region (E3), we measured a minor increase in collagen transcript density but a >3x increase in FB abundance. These results reveal a strong proliferative response of FBs localized to CM3-rich, high-fibrosis areas. FB proliferation thus accounted for most of the increase in FB-derived collagen transcripts in regions of high fibrosis. Meanwhile, for CMs, collagen transcript density was ∼5x increased in the high– *vs.* low-fibrosis regions (E3 *vs.* E1&E2), while abundance of CMs was similar across all microenvironments (Figure 4G). Taken together, these results show that local increases in FB-derived collagen transcripts are mainly caused by FB proliferation rather than transcriptional upregulation. By contrast, local increases in CM-derived collagen expression are entirely due to transcriptional activation.

To test if pro-fibrotic CM progression was associated with cell-type-specific gene expression signatures in local non-CM cells, we measured across all hexbins the correlation between CM3 abundance and gene expression from each non-CM cell type (Figure 4H). Our results reveal that CM3-associated FBs showed a shift toward a matrix-remodeling pro-fibrotic state, marked by increased Fn1 and Nox4 and a loss of the ECM– and TGF-β–regulatory components Ltbp4, Dcn, Fbln2, and Tgfbr3. These changes are consistent with a commitment of FBs to self-reinforcing myofibroblast activation. CM3-associated ICs showed decreased expression of Cd163 and Timp2, which indicates the presence of fewer anti-inflammatory (Cd163^+^) macrophages and presence of ICs with more aggressive matrix remodeling activity. CM3-associated ECs displayed signatures of a suppressed metabolic state, marked by downregulation of FA uptake/processing genes (Cd36, Lpl, Acsl1) together with reduced Pln and Ldha which might indicate endothelial dysfunction more broadly. Together, these findings reveal that the pro-fibrotic CM3 cardiomyocytes reside in a distinct microenvironment defined by cell-type-specific transcriptional changes in neighboring non-CM cells (Figure 4I).

### Predicting pro-fibrotic progression of cardiomyocytes from internal states of local non-cardiomyocytes

Based on our observation that pro-fibrotic CMs were found in distinct microenvironments, we reasoned that it may be possible to use the expression profiles of local non-CM cells to predict the progression of single CMs. We set out to do this with the help of a simple feed-forward neural net, which we hereafter refer to as Cell Health Inference Net (chiNet, written as χNet; Figure 5A).

**Figure 5.**
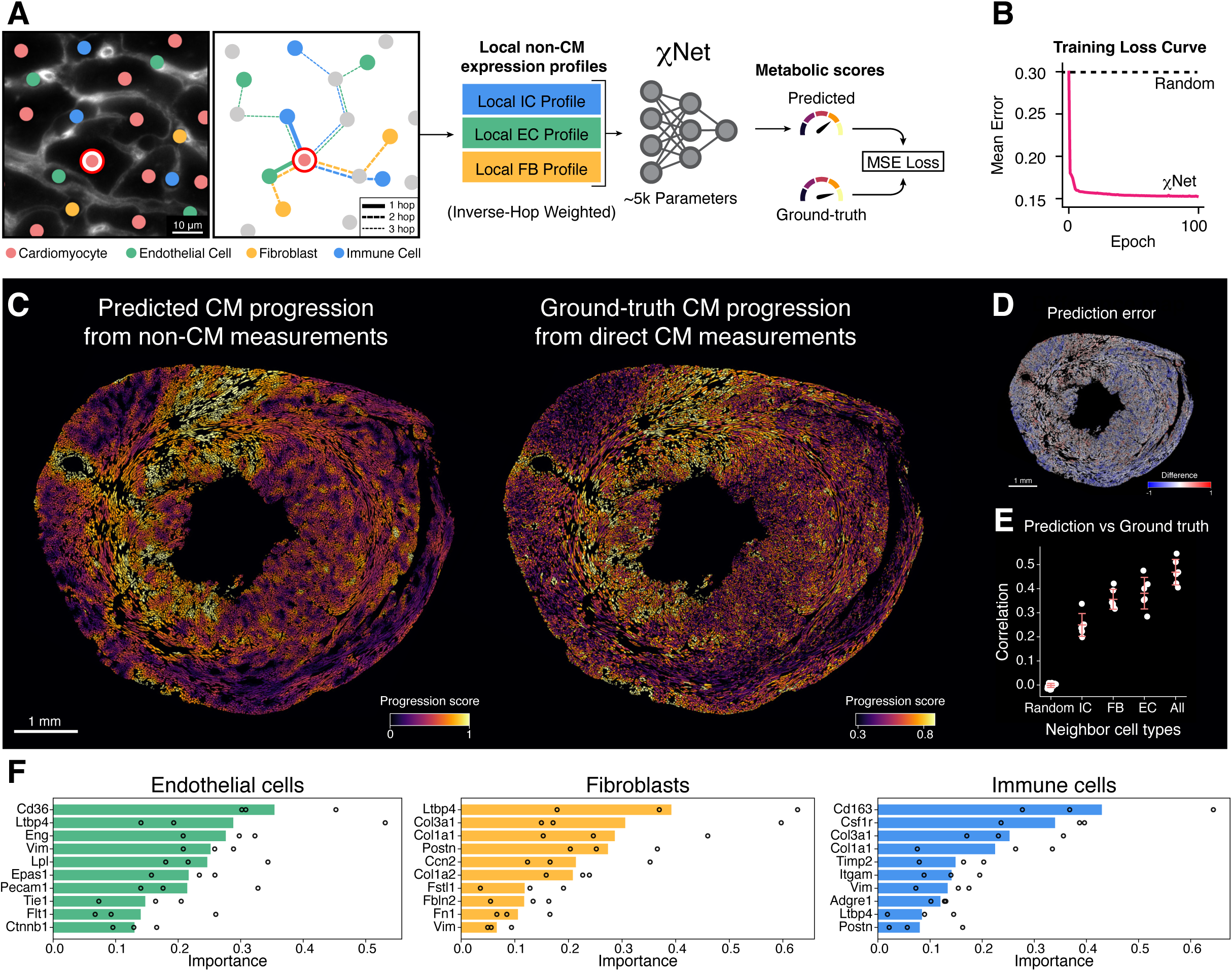
Reconstructing the spatial pattern of CM progression from nearby non-cardiomyocyte cells. (A) Design of χNet, a feed-forward neural net that uses the expression profiles of neighboring non-CM cells to predict single-cell CM progression scores in a TAC heart. Ground-truth CM progression scores were measured for each CM based on transcription of FA metabolism genes (Cd36, Acsl1, Acadm, Lpl), where decreased transcription of these genes corresponded to a greater progression score and *vice versa*. χNet-predicted CM progression scores were generated for each CM as follows: the expression profiles of neighboring non-CMs within a cutoff distance were distance-weighted (1/2/3 hop), combined within each non-CM cell type (IC, EC, FB), and used as input features. χNet uses these input features to predict CM progression score. (B) Performance of χNet on validation data consisting of a masked subset of the TAC heart section used for training. Performance was measured as mean absolute error relative to ground truth. Dashed line indicates the performance of a model with randomized model weights. (C) *Left:* χNet-prediction of CM progression scores in a TAC heart (TAC1), generated using transcription profiles of non-CM cells (up to 3-hop neighbors). This χNet model had been trained on a heart from a different mouse (TAC2). *Right:* Ground-truth CM progression scores for the same TAC heart section shown on the left. Ground-truth was determined from direct measurements of gene transcripts in CMs. The similarity of left and right images shows that model predictions are robust across animals, reflecting a conserved relationship between CM progression and the transcription profiles of non-CM cells in the local microenvironment. (D) Difference map showing differences between predicted progression scores and ground-truth. (E) Correlation between χNet predictions and ground truth across models trained using different subsets of neighbor non-CM cell types; “Random” shows performance when randomizing neighbors as a negative control. (F) Top informative genes differ across each non-CM cell types used for prediction. Importance of each gene for χNet predictions, as determined from the corresponding Shapley value (see Methods). Bars represent the mean importance across the three TAC heart sections and points correspond to individual heart sections.

We leveraged our cell connectivity graph to identify the cellular microenvironment for each CM (Figure 4A, Figure S4A), as well as the expression profiles of non-CM cells within these microenvironments. We then created a ground-truth dataset by measuring the severity of pro-fibrotic progression for every CM as a progression score normalized within each tissue from 0 to 1. To determine this progression score, we needed a reliable metric of the internal state of single CMs. Since the pro-fibrotic transcripts were expressed in high numbers only in the most severely progressed CMs, we were unable to use these transcripts to measure robust single-cell progression scores in every CM. Meanwhile, we had found that the decline in FA metabolism gene expression was tightly coupled to the increase in pro-fibrotic gene expression (Figure 2F, Figure 3, Figure S5A), reflecting a well-documented metabolic shift away from FA metabolism seen in CMs under stress^36^. Importantly, FA metabolism transcripts were highly abundant across all CMs, leading to a higher signal-to-noise ratio when measuring expression of these genes in single cells. We therefore used the aggregate expression of a co-expressed group of FA metabolism genes (Cd36, Acsl1, Acadm, Lpl) to more precisely determine the progression score of each CM (see Methods), and used these scores as a ground truth for the progression of CM state during pressure overload.

Discovery of relationships between cell state and/or other cell properties with the cellular microenvironment has previously been approached through a variety of different machine learning-based approaches^40–44^, as well as analytical methods^45^. Similar to χNet, many of these methods focus on inference of cellular gene expression^40–42^, while other approaches have been used to predict cell properties^43^, and for identification of environmental niches driving cell state and differentiation^44,46^. We designed χNet as a Multi-Layer Perceptron (MLP) to identify the most salient microenvironmental links to CM progression and to enable straightforward interpretability analysis.

We trained the χNet model to predict CM progression based on the transcription profile of neighboring non-CM cells. For training, we used one of our TAC datasets and set aside a portion of the dataset to allow for tracking model improvement during training. Model performance improved rapidly initially and then slowly converged toward a minimum error configuration that we chose for our final set of model weights (Figure 5B). Although we saw strong performance on our validation dataset, we worried that this performance could be the result of the model learning sample-specific patterns of disease. To test if the model had learned fundamental and more general associations between CM progression and local environment, we tested our model’s performance on a TAC tissue section from a different animal (that had not been used for training). For this test, we used only the new tissue’s cell connectivity graph and the expression profiles of non-CM cells. Using only this information, we then let the model predict the progression score of every CM in the previously unseen tissue section. The resulting χNet-predicted pattern of CM progression is shown in Figure 5C, *left image*. This pattern displays a striking resemblance to the pattern of ground-truth CM progression scores from the same animal (Figure 5C, *right image*). While the model tended to exaggerate the progression scores on both ends of the scale ( Figure 5D), it correctly reproduced the locations of nearly all areas with severe progression within the sample. To test the robustness of our findings, we repeated the training and testing procedure for every possible combination of our three TAC datasets, yielding a total of six experiments where in each experiment χNet was trained on a single tissue sample and evaluated on a sample from a different animal (Figure S5B). These experiments all showed a significant correlation between predicted and actual CM progression scores (Figure 5E, “*All*”, Figure S5C). Taken together, these results prove that in a pressure overloaded heart, the spatial patterns of CM progression are imprinted within the heart’s non-CM cells.

To determine which non-CM cell type was the most informative for predicting CM progression, we repeated these six experiments using only one non-CM cell type as an input for χNet. These experiments revealed that ECs and FBs were similarly informative, while ICs were consistently less informative (Figure 5E, “*IC*”, *FB*”, and “*EC*”, respectively). We speculate that the lower information content in ICs could be due to their relative sparsity within the tissue (see Figure 1F).

To estimate the distance over which non-CM cells could predict local CM progression, we repeated all prediction experiments using a variety of network attention distances. Specifically, we used as inputs the cell states of all non-CM cells within a distance ranging from 1 and up to 7 cells away (Figure S5D). Predictions improved when we increased the attention size from 1 to 5 cells but then reached a plateau. We speculate that the initial improved performance was due to the larger attention ranges capturing more cells, providing more measurements of local non-CM gene expression, thus leading to more accurate predictions. Meanwhile, the plateau could be due to an increasing trade-off between prediction accuracy and spatial precision, a limitation that could potentially be overcome by using a more sophisticated model architecture.

Finally, we asked which genes were the most informative within each of the non-CM cell types. To answer this question, we performed additional computational experiments to determine the average marginal contribution of each gene in each cell type towards the prediction of local CM progression (see Methods). Our results, shown in Figure 5F, reveal that different genes were the most informative in different cell types. In ECs, the expression of the FA carrier protein Cd36 was the most informative for predicting the progression of nearby CMs, suggesting that changes in FA transport by ECs might be linked to local CM progression. Meanwhile, in FBs, expression of the TGF-β regulator Ltbp4 was the most informative, which is consistent with the localized fibrosis seen in areas with more progressed CMs. Finally, in ICs the most informative gene was Cd163, suggesting that the presence of anti-inflammatory Cd163^+^ macrophages might play a role in modulating CM state. Many of these highly informative genes were also identified in our microenvironment analysis (Figure 4H), thus cross-validating our two different approaches. Taken together, our neural network analysis shows that under pressure overload, the spatial heterogeneity of CM progression is locally imprinted in the gene expression profiles of non-CM cells.

## Discussion

Here we measured single-cell geometries, spatial organization, physical connectivity, and transcription profiles in transverse sections of murine cardiac tissue, in healthy hearts and in hearts at early-stage pressure overload. Our ability to perform accurate spatial single-cell analysis in the densely crowded tissue of the heart crucially relied on Cellouette, a new cell segmentation approach that improved coverage and accuracy compared to previous approaches. The high coverage of Cellouette ensured minimal loss of cells and transcripts in population and microenvironment analyses. The high segmentation accuracy ensured that our discoveries were not caused by misattributed transcripts, a common occurrence in other spatial studies^47^. Based on the improvement in spatial analysis capabilities enabled by Cellouette in the heart, we propose that similarly improved cell segmentation in other crowded tissues will unlock the potential of spatial transcriptomics to reveal relationships between cells and their environment. Importantly, we found that accurate image-based cell segmentation relied in equal parts on the use of appropriate machine learning models and on the use of an unambiguous cell boundary stain, the latter of which will need to be optimized in each tissue of interest.

We discovered that pressure overload causes CMs to adopt a pro-fibrotic transcriptional profile that included substantial expression of several fibrosis-related genes (eg. Col1a1, Col1a2, Postn) previously attributed mainly to FBs. Since the spatial pattern of fibrosis-related gene expression in CMs coincided with the pattern of tissue fibrosis, it is tempting to speculate that the expression of these genes by CMs play a functional role in fibrosis. However, a previous study showed that CM-specific ablation of the collagen chaperone Hsp47 in a TAC model did not lead to significant reduction of tissue fibrosis at a late timepoint, four weeks post-surgery^48^, strongly suggesting that CMs are not a major source of ECM production, and leaving the functional role of CM expression of fibrillar collagen genes a bit of a mystery. Our study contributes to this picture in several ways. First, we found that local increases in FB-derived collagen transcript counts were largely driven by FB proliferation rather than intracellular upregulation (Figure 4F). Since the CM-derived increase in collagen transcripts was entirely due to transcriptional upregulation (Figure 4G), we speculate that the CM response may be faster than the FB response, potentially allowing for fine-tuned adjustment of ECM composition over shorter time scales. Second, since we found that expression of fibrosis-related genes coincided with a shift in expression of genes involved in FA metabolism and calcium handling, we speculate that this pro-fibrotic transcriptional reprogramming reflects a broader cardiomyocyte state transition and may contribute to disease progression through mechanisms independent of direct collagen secretion^9^.

In the pressure-overloaded heart, we identified novel relationships linking the coupled metabolic and pro-fibrotic progression of a CM to the cellular and molecular composition of its local microenvironment. While our current findings do not distinguish between relationships that are correlated or causal, we suggest that the specific genes we identified and the cells in which they are expressed can serve as a candidate list for cell type-specific gene programs that may impact CM progression under pressure overload. More broadly, we anticipate that our approach to find candidate microenvironmental factors can be combined with multiplexed cell-specific perturbations^49^ to identify microenvironmental factors that drive CM resilience during pathological stress and thus can present therapeutic targets for prevention or treatment of heart disease.

We foresee that our resource will enable discovery of new relationships between cell state, shape, and environment. For instance, we found a non-linear relationship between CM size and Acta1 expression in the pressure overloaded hearts (Figure 2G). Using a similar approach, our data could be used to discover drivers and modulators of CM hypertrophy *in situ*.

We showed that CM shape can be used to determine muscle fiber orientation in a single tissue section (Figure 1I), which will enable studies into how the orientation of a muscle fiber affects its resilience under mechanical stress. The combination of single-CM orientation and transcriptional state could also inform multiscale biomechanical models that integrate clinical measurements, including blood pressure and ventricular geometry, to estimate local wall stress and predict CM physiological responses^32,50^.

Our discovery that CM progression is imprinted in nearby non-CM cells opens the possibility of *in vivo* “remote sensing” of CM state. CMs are notoriously difficult to examine *in vivo*, however we found that shifts in the internal states of non-CM cells are sufficient for predicting the state of adjacent CMs. We envision that this discovery can enable determination of CM health *in vivo* by examining more accessible non-CM cells. For instance, minimally invasive approaches could be used to assess the state of cells on the surface of the heart or the cells lining the vasculature of major coronary arteries. A computational model similar to χNet could then be used to infer the health of the less accessible CMs located nearby, and thereby enable more accurate diagnostics of cardiac health, injury, and disease.

## Author contributions

P.K. and R.T.L. conceived the study. A.V., Y.L., and R.T.L. designed the gene panel. Y.L. designed and performed MERFISH experiments. K.M.H., U.M., and T.D.P. supervised A.S. and V.V.T. during the development of segmentation methods for fluorescence image data. A.S., V.V.T., A.M.C., Y.L., and J.W. developed Cellouette and performed cell segmentation. Y.L. and D.G. performed cell classification. K.C. and J.F. designed and performed SVG analysis. A.M.C. and Y.L. designed and performed network analysis. Y.L., A.M.C., and J.W. designed and performed microenvironment analysis. J.C. designed and performed cell orientation and χNet analysis. Y.L., A.M.C., K.C., and J.C. conducted data visualization and figure preparation. Y.L., P.K., and A.M.C. wrote the manuscript with input from all authors. P.K. supervised the study.

## Acknowledgments

This work was supported by a Beckman Young Investigator Award (P.K.), a W.M. Keck Foundation Award (P.K.), a Pioneer Fund postdoctoral fellowship (Y.L.), a Salk Women & Science research award (Y.L.), an Edwards-Yeckel Postdoc Professional Development Award (Y.L.), NIH grant R01 HL169291 (R.T.L.), NIH grant R35 GM142889 (K.C., J.F.), NIH Grant R56 MH139176 (K.M.H.), NSF Grants 1707356, 2014862, and 2219894 (K.M.H.). We would like to acknowledge contributions by Bojing Blair Jia, Leonardo Sepúlveda, Won Jun, Julia Rune, Patrick Sun, Raghu Parupudi, Deepak Pant, and Jenny Weixiu Dong.

