## Supplemental Figures S1 – S5 for "Spatial imprints of emergent cardiomyocyte states in the pressure-overloaded heart"

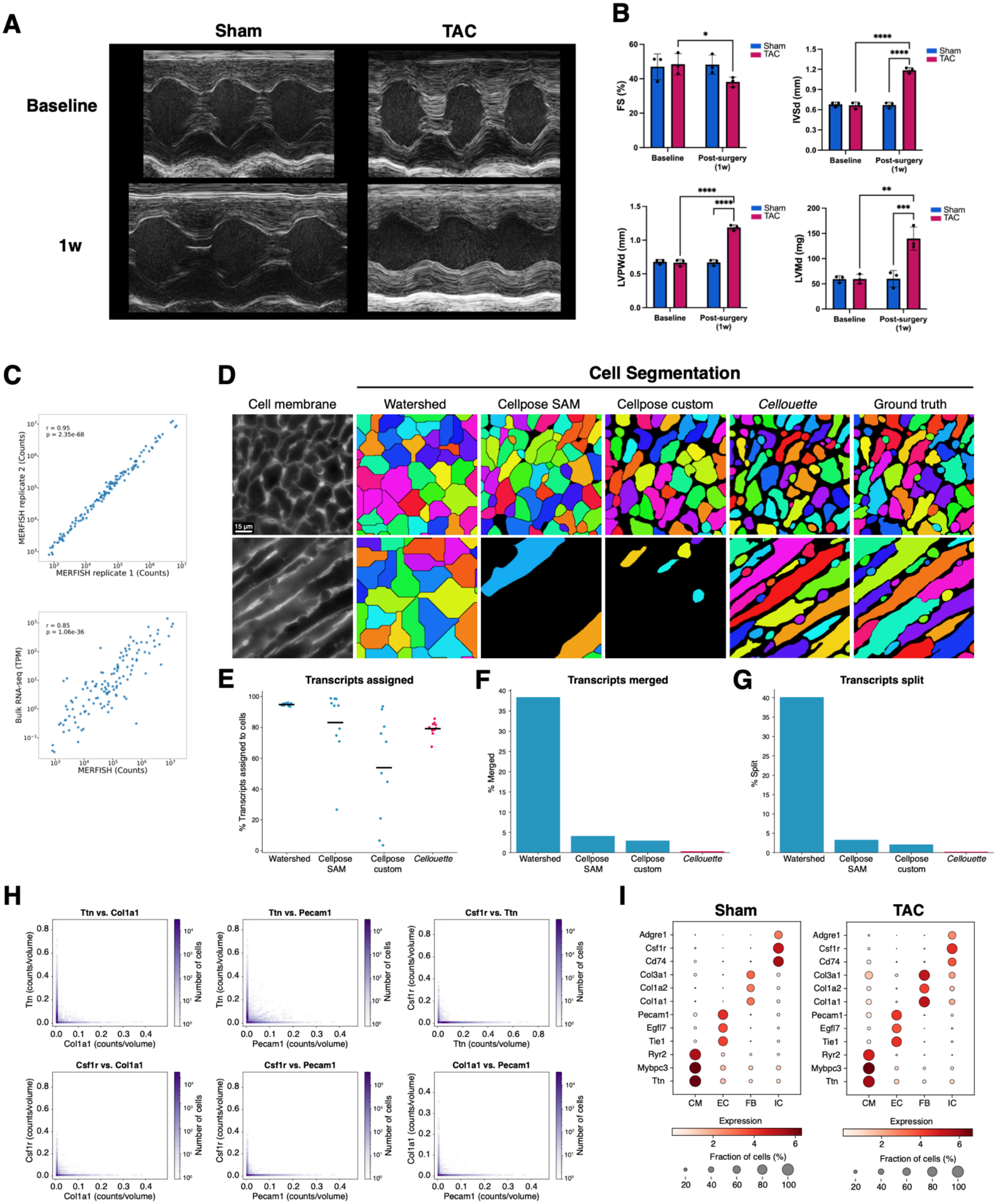
A
Sham
TAC
Baseline
1w
B
FS (%)
IVSd (mm)
LVPWd (mm)
LVMd (mg)
Sham
TAC
Baseline
Post-surgery (1w)
C
r = 0.95
p = 2.35e-68
MERFISH replicate 2 (Counts)
MERFISH replicate 1 (Counts)
r = 0.85
p = 1.06e-36
Bulk RNA-seq (TPM)
MERFISH (Counts)
D
Cell Segmentation
Cell membrane
Watershed
Cellpose SAM
Cellpose custom
Cellouette
Ground truth
15 µm
E
Transcripts assigned
% Transcripts assigned to cells
F
Transcripts merged
% Merged
G
Transcripts split
% Split
Watershed
Cellpose SAM
Cellpose custom
Cellouette
H
Ttn vs. Col1a1
Ttn vs. Pecam1
Csf1r vs. Ttn
Csf1r vs. Col1a1
Csf1r vs. Pecam1
Col1a1 vs. Pecam1
Ttn (counts/volume)
Csf1r (counts/volume)
Col1a1 (counts/volume)
Pecam1 (counts/volume)
Number of cells
I
Sham
TAC
Adgre1
Csf1r
Cd74
Col3a1
Col1a2
Col1a1
Pecam1
Egfl7
Tie1
Ryr2
Mybpc3
Ttn
CM
EC
FB
IC
Expression
Fraction of cells (%)

**Figure S1. Pressure overload physiology and quality control for single-cell analysis.**
(A) Echocardiographic validation of early-stage cardiac remodeling following transverse aortic constriction (TAC)**.** Representative M-mode echocardiographic images of sham-operated and TAC hearts at baseline (prior to surgery) and one-week post-surgery.

(B) Quantitative echocardiographic analysis showing fractional shortening (FS), interventricular septal thickness in diastole (IVSd), left ventricular posterior wall thickness in diastole (LVPWd), and left ventricular mass in diastole (LVMd) at baseline and one-week post-surgery. At baseline, no significant differences were observed between sham (n=3) and TAC (n=3) groups. One week after surgery, TAC mice exhibited significantly reduced FS and significantly increased IVSd, LVPWd, and LVMd, compared to sham-operated controls, confirming effective induction of pressure overload and ventricular hypertrophy. Data are presented as mean ± s.d.; statistical analysis was performed using two-way ANOVA followed by Tukey's multiple comparisons test; *$p < 0.05$, **$p < 0.01$, ***$p < 0.001$, ****$p < 0.0001$.

(C) Reproducibility and validation of MERFISH transcript measurements. Top: correlation between average transcript counts per gene determined by MERFISH and gene expression levels (TPM) determined by bulk RNA-seq from mouse heart (Tabula Muris Senis dataset; Almanzar *et al.*, *Nature* 2020). Pearson correlation coefficient (r) = 0.85; p = 1.06e-36. Axes are displayed on log scale. Bottom: correlation of average transcript counts per gene between two independent MERFISH biological replicates from mouse heart sections. Pearson correlation coefficient (r) = 0.95; p = 2.35e-68. Axes are displayed on log scale.

(D) Comparison of cell segmentation methods in regions of the ventricular heart sections with cross-sectioned CMs (*top row*) and longitudinally sectioned CMs (*bottom row*). DAPI-based watershed segmentation, which relies on nuclear detection, failed to accurately segment CMs, especially those whose nuclei were not captured within the tissue section. The Cellpose models performed well on cross-sectioned CMs and smaller non-CM cells but failed to properly segment longitudinally oriented CMs. Cellouette showed robust segmentation performance, successfully identifying both cross-sectional and longitudinal cardiomyocytes, as well as small non-CM cells, across diverse and crowded myocardial environments.

(E) Percent of transcripts partitioned in the ground-truth test data that were partitioned into cells using different segmentation approaches. Watershed showed nearly complete (but incorrect) partitioning. Cellpose approaches showed variable performance across different regions of the sample, leading to CM orientation bias. Cellouette showed consistent performance across the test data set.

(F) Transcripts merged into a single cell that were part of multiple cells in the ground-truth data. Values given as percent of total partitioned transcripts in the ground-truth data.

(G) Transcripts split into multiple cells that were found in a single cell in the ground-truth data. Values given as percent of total partitioned transcripts in the ground-truth data.

(H) Validation of cell segmentation and transcript partitioning accuracy using mutually exclusive expression of cell-type-specific marker genes. Two-dimensional histograms showing co-expression of marker genes from different cell types across all segmented cells from three sham samples. The observed mutual exclusivity of marker gene expression indicates a low level of cross-contamination of transcripts between neighboring cells. Marker genes were selected for each major cell type: Csf1r (immune cells) vs. Col1a1 (fibroblasts), Csf1r (immune cells) vs. Pecam1 (endothelial cells), Ttn (cardiomyocytes) vs. Col1a1 (fibroblasts), and Ttn (cardiomyocytes) vs. Pecam1 (endothelial cells). Color intensity represents the number of cells in each bin.

(I) Cell-type-specific expression patterns in sham and TAC hearts. Dot plots showing the expression of representative marker genes across major cardiac cell types: CM, EC, FB, and IC in sham (left) and TAC (right) hearts. Dot color intensity represents average gene expression, and dot size represents the fraction of cells expressing each gene within a given cell type.

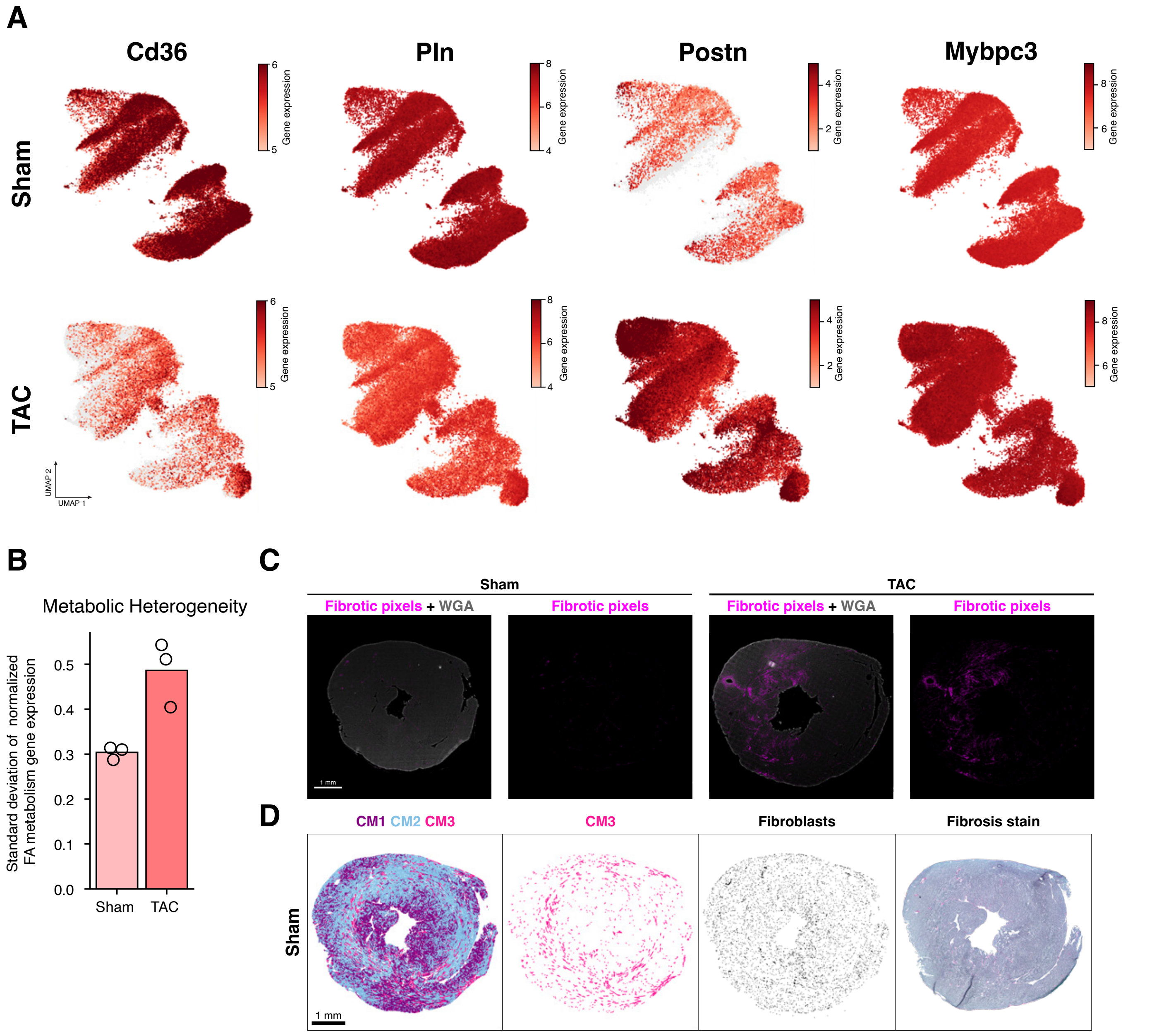


**Figure S2. CM transcriptional differences and spatial patterns of fibrosis in sham and TAC hearts.** (A) Expression pattern of selected genes across cardiomyocytes in sham and TAC hearts. UMAP visualizations showing the expression of representative genes involved in fatty acid metabolism (Cd36), calcium handling (Pln), fibroblast activation (Postn), and sarcomeric structure (Mybpc3) in sham (top row) and TAC (bottom row) cardiomyocytes. Color intensity indicates normalized gene expression levels in individual cells.

(B) TAC hearts exhibit increased metabolic heterogeneity. Standard deviation of expression of normalized FA metabolism genes (Cd36, Acsl1, Acadm, Lpl) across cardiomyocytes of sham and TAC hearts. Bars indicate the group mean; points represent individual hearts.

(C) Representative spatial distribution of fibrosis in sham and TAC hearts. Whole midventricular cross-sections from sham (*left*) and TAC (*right*) hearts stained with WGA (*gray*) to highlight cell boundaries, overlaid with computationally identified fibrotic pixels (*magenta*) extracted from fibrosis histology stain (Sirius red) of an adjacent tissue section. Scale bar, 1 mm.

(D) Spatial distribution of CM states in a sham heart; location of pro-fibrotic CMs (CM3); location of FBs; fibrosis histology stain (Sirius red) of an adjacent tissue section.

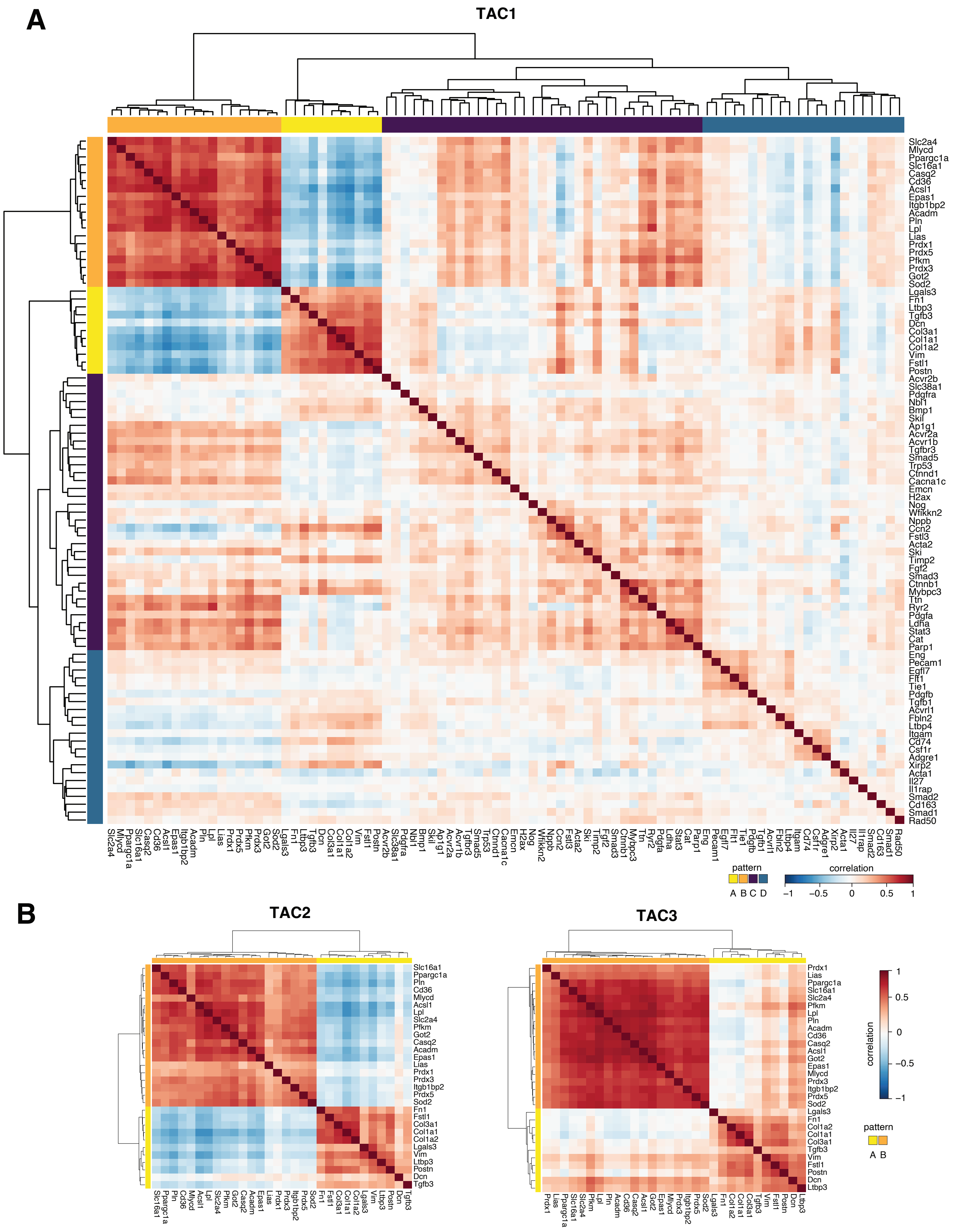
A
TAC1
Slc2a4
Mlycd
Ppargc1a
Slc16a1
Casq2
Cd36
Acsl1
Epas1
Itgb1bp2
Acadm
Pln
Lpl
Lias
Prdx1
Prdx5
Pfkm
Prdx3
Got2
Sod2
Lgals3
Fn1
Ltbp3
Tgfb3
Dcn
Col3a1
Col1a1
Col1a2
Vim
Fstl1
Postn
Acvr2b
Slc38a1
Pdgfra
Nbl1
Bmp1
Skil
Ap1g1
Acvr2a
Acvr1b
Tgfbr3
Smad5
Trp53
Ctnnd1
Cacna1c
Emcn
H2ax
Nog
Wfikkn2
Nppb
Ccn2
Fstl3
Acta2
Ski
Timp2
Fgf2
Smad3
Ctnnb1
Mybpc3
Ttn
Ryr2
Pdgfa
Ldha
Stat3
Cat
Parp1
Eng
Pecam1
Egfl7
Flt1
Tie1
Pdgfb
Tgfb1
Acvrl1
Fbln2
Ltbp4
Itgam
Cd74
Csf1r
Adgre1
Xirp2
Acta1
Il27
Il1rap
Smad2
Cd163
Smad1
Rad50
pattern
A B C D
correlation
−1
−0.5
0
0.5
1
B
TAC2
Slc16a1
Ppargc1a
Pln
Cd36
Mlycd
Acsl1
Lpl
Slc2a4
Pfkm
Got2
Casq2
Acadm
Epas1
Lias
Prdx1
Prdx3
Itgb1bp2
Prdx5
Sod2
Fn1
Fstl1
Col3a1
Col1a1
Col1a2
Lgals3
Vim
Ltbp3
Postn
Dcn
Tgfb3
TAC3
Prdx1
Lias
Ppargc1a
Slc16a1
Slc2a4
Pfkm
Lpl
Pln
Acadm
Cd36
Casq2
Acsl1
Got2
Epas1
Mlycd
Prdx3
Itgb1bp2
Prdx5
Sod2
Lgals3
Fn1
Col1a2
Col1a1
Col3a1
Tgfb3
Vim
Fstl1
Postn
Dcn
Ltbp3
correlation
1
0.5
0
−0.5
−1
pattern
A B

**Figure S3. CM-specific Spatially Variable Gene (SVG) patterns in different TAC hearts.**
(A) Hierarchically clustered correlation matrix of spatially binned gene expression for all genes in TAC1 with a CM expression level above background.

(B) Correlation matrix for TAC2 and TAC3 of spatially binned gene expression for the genes identified as part of the top two clusters in TAC1.

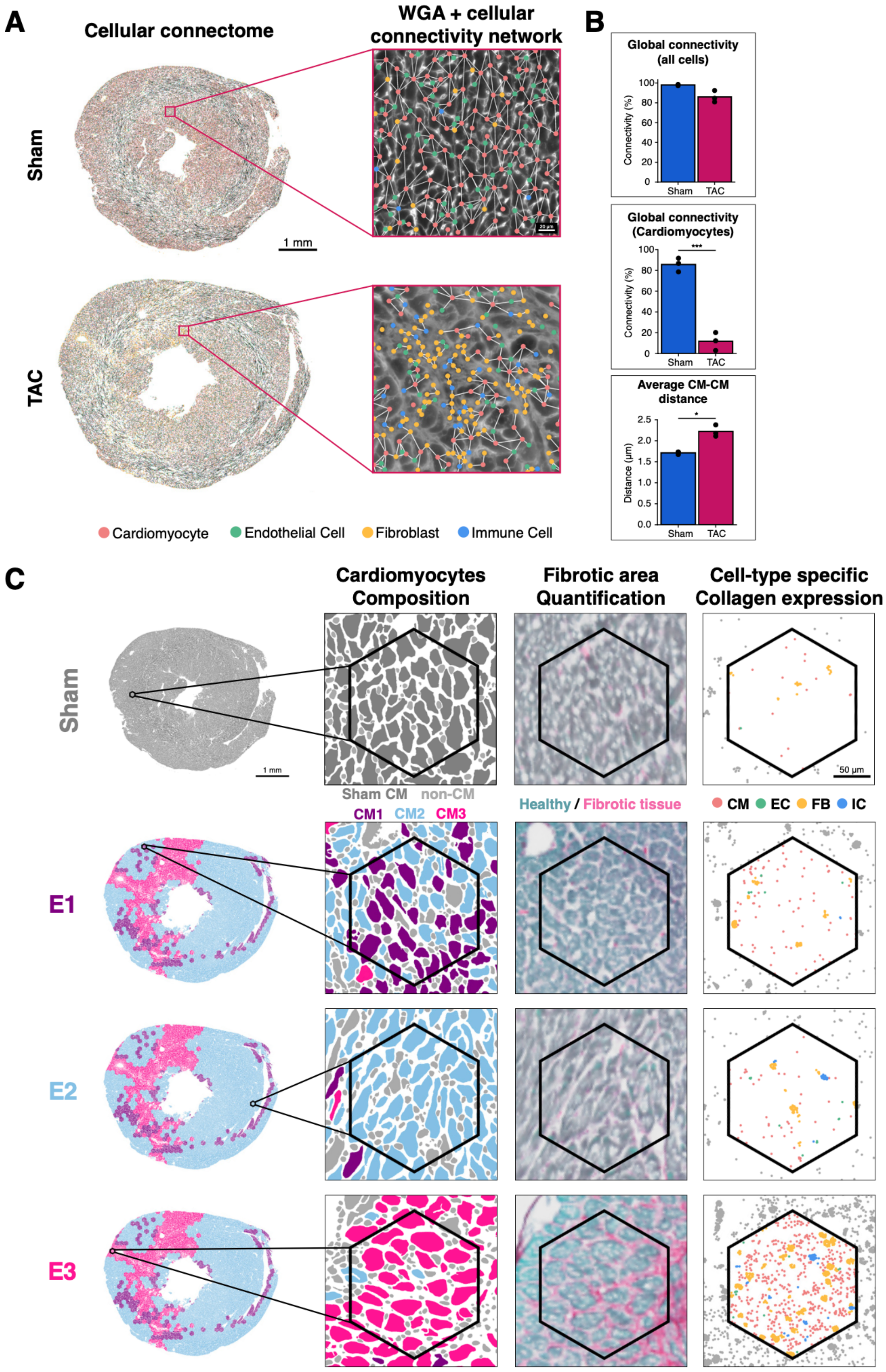

A
Cellular connectome
WGA + cellular connectivity network
Sham
TAC
1 mm
Cardiomyocyte
Endothelial Cell
Fibroblast
Immune Cell
B
Global connectivity (all cells)
Connectivity (%)
Sham
TAC
Global connectivity (Cardiomyocytes)
***
Average CM-CM distance
*
Distance (µm)
C
Cardiomyocytes Composition
Fibrotic area Quantification
Cell-type specific Collagen expression
Sham CM
non-CM
CM1
CM2
CM3
Healthy / Fibrotic tissue
CM
EC
FB
IC
50 µm
E1
E2
E3

**Figure S4**. **Cell connectivity graphs and hexbin analysis reveal distinct cellular environments in sham and TAC hearts.**
(A) Spatially resolved network graphs illustrating cellular interactions among cardiomyocytes (CMs, red), fibroblasts (FBs, yellow), endothelial cells (ECs, green), and immune cells (ICs, blue) in representative sham (top) and TAC (bottom) heart sections. Insets highlight diverse cellular interactions, with edges indicating physical proximity (threshold ≤ 3 μm).

(B) Quantification of cellular connectivity and distances between neighboring cells: Top: Global connectivity measured as the percentage of cells within the largest interconnected network graph. Sham hearts exhibit high connectivity, while TAC hearts show slightly reduced connectivity among all cell types. Middle: Global connectivity among cardiomyocytes, significantly reduced in TAC compared to sham hearts. Mean ± s.d., ***$p < 0.001$ (unpaired two-tailed t-test). Bottom: Average shortest distance between neighboring cardiomyocytes, measured using network graphs constructed with a 10 μm threshold to connect all nodes while quantifying only cardiomyocyte–cardiomyocyte distances. Inter-cardiomyocyte distances significantly increased in TAC hearts. Mean ± s.d., *$p < 0.05$ (unpaired two-tailed t-test).

(C) CM microenvironment analysis pipeline. Single-cell maps and Sirius Red histology images from sham and TAC heart sections were computationally aligned and rasterized into hexagonal bins “hexbins”, each representing a local cardiomyocyte microenvironment. Each hexbin was classified as E1, E2, or E3 corresponding to the most enriched cardiomyocyte state (CM1, CM2, or CM3, respectively). The sham microenvironment was represented by randomly selected hexbin from a sham heart. For each hexbin, microenvironmental features were quantified, including the composition of CM states, the percentage of area covered by fibrosis, and cell-type-specific expression of collagen genes.

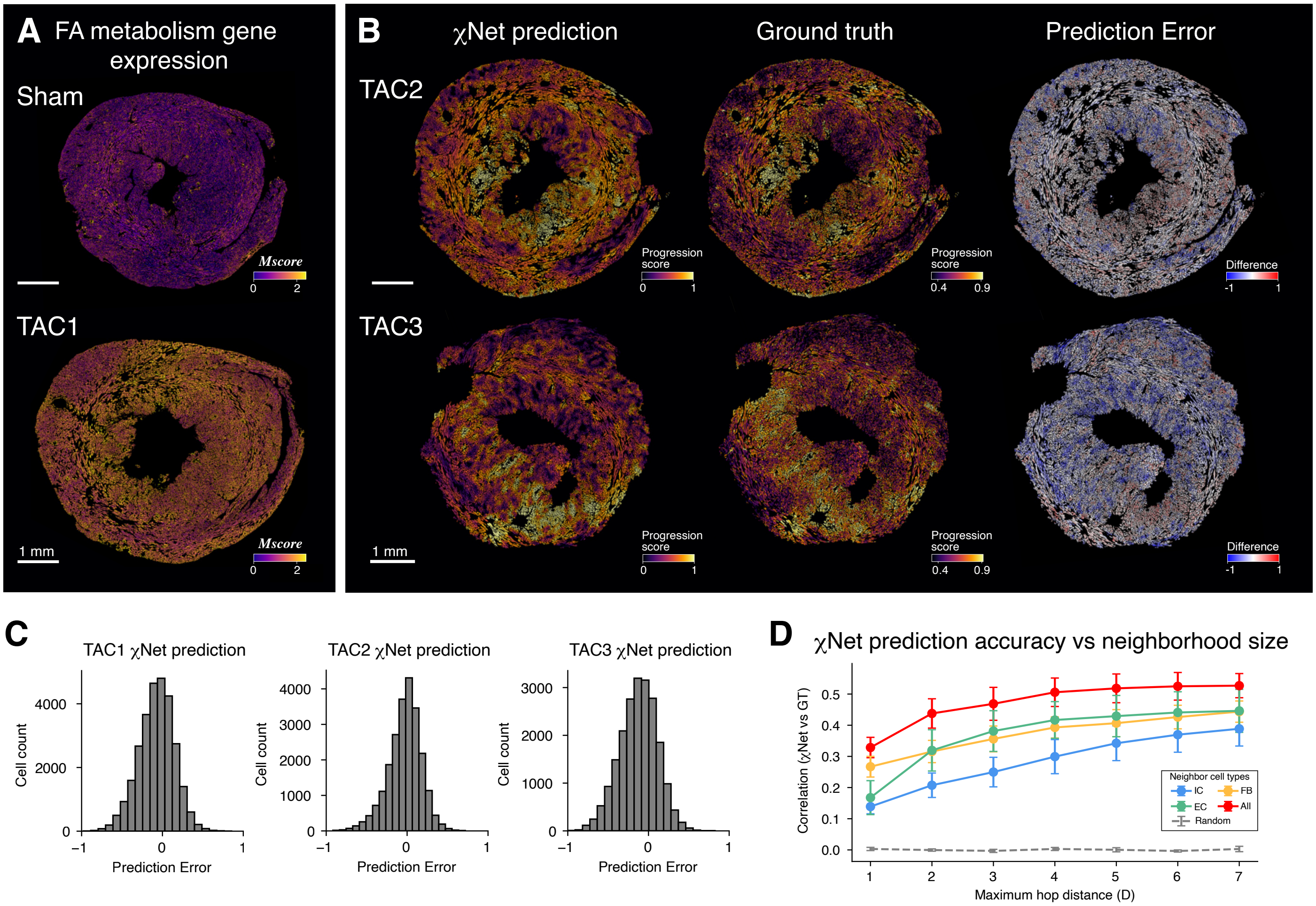


**Figure S5. CM progression patterns and χNet performance across different animals.**
(A) Spatial maps of CMs in sham and TAC heart sections, showing FA metabolism gene expression on an inverted scale (yellow indicates lower expression, corresponding to greater progression). These maps reveal that the TAC heart CMs display greater heterogeneity and greater overall progression. To correct for batch effects, we leveraged the fact that Mybpc3 was found to be both highly expressed in all CMs and expressed at a stable level across conditions and CM states (see for instance Figures 2F, S1I & S2A). For each tissue section, batch-corrected single-cell expression values were calculated by normalizing single-cell expression values to the section's mean Mybpc3 expression. We then computed a per-cell aggregate FA metabolism expression value $M_i$ by summing batch-corrected expression values across the FA metabolism genes Cd36, Acsl1, Acadm, and Lpl. We then inverted the scale to make lower FA metabolism expression correspond to a higher score: $Mscore_i = max_j(M_j) - M_i$

(B) χNet generalizes across animals. For TAC2 and TAC3 hearts, ground-truths (*left*) are compared with χNet predictions (*center*) generated by a model trained on a different TAC sample, demonstrating χNet's ability to capture the major spatial features of CM metabolic shift across animals. Difference maps (*right*) show difference between ground truth and prediction.

(C) Quantitative evaluation of χNet performance. For each individual CM cell, prediction error was defined as the difference between its ground-truth and χNet-predicted metabolic progression score. Histograms show symmetric errors and consistent distributions across TAC1, TAC2, and TAC3.

(D) Prediction accuracy (correlation between χNet prediction and ground truth) as a function of network size (neighbor degree n). Accuracy improves as more neighborhood layers are included and is highest when all non-CM cell types are used. Randomized-neighbor controls (*gray*) show no predictive power.
