## Supplementary material for "Spatial imprints of emergent cardiomyocyte states in the pressure-overloaded heart": Methods

#### 1. Animals

**Transverse Aortic Constriction (TAC) procedure**

8-12-week-old adult male mice (C57BL/6; The Jackson Laboratory) were anesthetized with ketamine (50 mg/kg, Dechra Veterinary, 07-894-8462) and xylazine (5 mg/kg, VetOne, 1XYL004) via intraperitoneal injection for initial induction, followed by isoflurane (0.75–1.5%) for complete anesthesia. The chest cavity was entered through the second intercostal space at the left upper sternal border, and the transverse aorta was isolated between the carotid arteries. In TAC mice, aortic constriction was performed by tying a 7-0 silk suture ligature around the aorta against a 27–27.5-gauge needle (selected based on body weight), which was promptly removed to produce a constriction of approximately 0.4 mm in diameter. After the procedure, the chest was closed with 6-0 silk sutures. Buprenorphine (0.1 mg/kg, 100 µL per mouse, McKesson, 3475027) was administered 15–30 min prior to anticipated recovery. Sham mice underwent the same surgical procedure except for the suture ligation.

**Cardiac physiology measurements using transthoracic echocardiography**

Prior to echocardiography, a depilatory cream was applied to the anterior chest wall to remove hair. Mice were anesthetized with 5% isoflurane for 15 seconds and then maintained at 0.5% throughout the echocardiographic examination. Small needle electrodes were inserted into one upper and one lower limb for simultaneous electrocardiogram recording. Transthoracic echocardiography (M-mode and 2-dimensional) was performed using a high-resolution ultrasound system with a 32–55 MHz linear transducer (FUJIFILM VisualSonics Vevo 2100). From these readings, measurements of chamber dimensions and wall thicknesses were obtained, and left ventricular fractional shortening and circumferential shortening velocity were calculated. Percentage fractional shortening (%FS) was used as an indicator of systolic cardiac function.

#### 2. Cardiac MERFISH data collection

**Heart tissue collection**

Mice were euthanized using $CO_2$, and the chest cavity was opened to expose the heart. The right atrium was cut with scissors, and the heart was slowly perfused from the apex with 10 mL of ice-cold PBS containing Murine RNase inhibitor (1:5000, New England Biolabs, M0314L) using a 26G needle. The heart was then quickly extracted, rinsed in ice-cold 1XPBS, and excess 1XPBS was removed with Kimwipe. The heart was horizontally bisected at the midventricular position and embedded in a cryomold filled with optimal cutting temperature (O.C.T.) compound (Tissue-Tek O.C.T, 4583), immediately frozen using a dry ice and ethanol bath, and stored at –80 °C until cryosectioning.

**Heart tissue cryosection**

Frozen hearts were cryosectioned at –20 °C using a cryostat (Leica CM1950). Trimmed sections were discarded until both the left and right ventricular regions were reached. Two consecutive 10-µm-thick sections were then collected: one was placed on a pre-processed 40-mm coverslip for MERFISH experiments, and the other was mounted on a microscope slide for histological staining. Coverslips were prepared as previously described[1]. Briefly, 40-mm-diameter #1.5 coverslips (Bioptechs, 40-1313-03193) were cleaned in a 1:1 mixture of 37% (vol/vol) Hydrochloric Acid (Fisher Scientific, A144-500) and methanol (Fisher Scientific, AA22909K7) at room temperature for 30 min. They were then washed three times with MilliQ water and once with 70% ethanol (vol/vol) (Fisher Scientific, BP2818500), followed by drying with nitrogen gas. Dried coverslips were incubated in a solution containing 0.1% (vol/vol) triethylamine (Millipore Sigma, TX1200) and 0.2% (vol/vol) allyltrichlorosilane (Millipore Sigma, 107778) in chloroform (Fisher Scientific, AAJ67241AP) for 30 min at room temperature. After incubation, coverslips were washed once with chloroform and once with ethanol, then baked in a 60 °C oven for 1 h to dehydrate and stabilize the silane layer. Silanized coverslips were stored at room temperature in a desiccated chamber.

Silanized 40-mm coverslips were further processed before cryosectioning. Briefly, each coverslip was incubated with poly-D-lysine solution (0.1 mg/mL, Santa Cruz Biotechnology, sc-136156) at room temperature for 1 hour. Coverslips were incubated with orange fluorescent fiducial beads (1:10000 dilution in 1× PBS, Thermo Fisher Scientific, F8800) for 10 minutes at room temperature to allow bead attachment. Following bead attachment, coverslips were fixed with 3 mL of 4% freshly diluted paraformaldehyde (Electron Microscopy Sciences, 15714) for 10 minutes at room temperature, washed three times with 1XPBS and dried using nitrogen gas.

**MERFISH library design and assembly**

To identify transcriptionally distinct cell populations in the healthy and pressure-overloaded mouse heart, a panel of 128 genes was designed. Among the 128 genes, known cell type marker genes were included for cardiomyocytes, fibroblasts, endothelial cells, and immune cells, as well as genes involved in hypertrophy, fibrosis, inflammation, and metabolism. Each of the 128 genes was assigned a unique binary barcode drawn from a 16-bit, Hamming-Distance-4, Hamming-Weight-4 barcode scheme. Barcodes were separated from one another by a Hamming distance of at least 4, meaning that a minimum of four bits would need to be misread to convert one valid barcode into another. A constant Hamming weight of 4 (i.e., the number of “1” bits per barcode) was used to avoid measurement bias due to asymmetric error rates in “1” to “0” and “0” to “1” transitions. The full codebook is listed in Supplemental Table S2. An additional 12 barcodes were included as ‘blank’ barcodes, unassigned to any gene, to measure the false-positive detection rate in MERFISH decoding. MERFISH encoding probes targeting the mRNA transcripts from the 128 genes were designed as previously described[2]. Each MERFISH encoding probe was designed to contain: (1) a 30-mer sequence complementary to the selected gene’s mRNA transcript; (2) three 20-mer readout sequences matching the on-bits of the gene’s barcode; and (3) two amplification priming regions, a 20-nt primer binding site at the 5′ end and the reverse complement of the T7 promoter at the 3′ end, as previously described[3]. To design the transcript-targeting regions of the encoding probes, first all possible 30-mer target sequences within each gene transcript were identified as previously described[2]. Briefly, 90–100 target sequences were randomly selected with the following criteria: GC content of 43%–63%, melting temperature ($T_m$) between 66 and 76 °C, a gene-specificity index between 0.75 and 1, an isoform-specificity index between 0.75 and 1, and no homology greater than 15 nt to rRNAs or tRNAs. For the readout sequences on the encoding probes, each of the 16 bits in the MERFISH barcode was assigned a 20 nt, three-letter nucleic acid sequence corresponding to the four ‘1’ on-bits for each gene. The encoding probe sequences for all 128 genes are listed in Supplemental Table S4. The encoding probe library was purchased as an oligo pool (Twist Bioscience) and amplified as previously described[2]. Briefly, the oligonucleotide pool was PCR-amplified to generate *in vitro* transcription templates (Phusion Hot Start Flex 2X Master Mix, New England Biolabs, M0536L) using:
Forward primer: 5′-TTGGGCGTGCACGTCAATTC-3′
Reverse primer: 5′-TAATACGACTCACTATAGGGCCATTGCCCGCGAGGTCGAG-3′
The amplified pool was then transcribed into RNA (HiScribe™ T7 Quick High Yield RNA Synthesis Kit, NEW ENGLAND BIOLABS, E2050S), and reverse-transcribed into single stranded DNA (Maxima H Minus Reverse Transcriptase, Thermo Fisher Scientific, EP0753) using:
Forward primer: 5′-TTGGGCGTGCACGTCAATrUC–3′
Reverse primer: 5′-TAATACGACTCACTATAGGGCCATTGCCCGCGAGGTCGAG-3′
RNA was then removed via alkaline hydrolysis, and DNA was purified using custom made magnetic nucleic acid purification beads. Final single stranded DNA probes were resuspended in RNase-free water and stored at −20 °C.

**WGA-oligonucleotide conjugation for cell boundary staining**

Wheat Germ Agglutinin (WGA; Vector Labs, L-1020-10) was conjugated to a 5′-Acrydite and 3′-Azide–modified oligonucleotide via a two-step copper-free click chemistry protocol. First, WGA (1 mg/mL in 25 mM HEPES-buffered saline, pH 7.5) was incubated with DBCO-PEG5-NHS ester (5 mM in DMSO; Kerafast) at room temperature. After the DBCO reaction, excess linker was removed by centrifugal filtration (Millipore Sigma, Amicon® Ultra 10 kDa MWCO filters, UFC201024) at 14,000 × g for 10 min, followed by buffer exchange into fresh HEPES-buffered saline. Next, the DBCO-modified WGA was reacted with the azide-modified oligonucleotide (sequence: 5′-Acrydite-GGGTAGTGGGAATGATTTAT-3′-Azide, 100 µM in HEPES-buffered saline, IDT). The reaction proceeded at 4°C overnight for at least 16 h. Unreacted oligonucleotide was removed by centrifugal filtration, and the WGA–oligo conjugate was stored at −20 °C until use.

**Design and construction of MERFISH readout probes**

For the 140-gene panel used in this study, 16 readout probes were designed as previously described[3], and ordered synthesized and purified (Eurofin Genomics). Each of these probes was complementary to one of the 16 readout sequences corresponding to one of the 16 bits in the barcodes. Each readout probe was conjugated to one of the two dye molecules Alexa750 or Cy5 via a disulfide linkage. Upon arrival from the vendor, these readout probes were resuspended immediately in Tris-EDTA (TE) buffer, pH 8 (Thermo Fisher, BP2473), to a concentration of 100 µM and stored at −20 °C. The readout probe sequences are listed in Supplemental Table S3.

**Preparation of cardiac tissue sections for MERFISH imaging**

After sectioning and placing the 10 µm-thick tissue section on a coverslip, each section was immediately fixed with 4% freshly diluted paraformaldehyde (Electron Microscopy Sciences, 15714) in 1x PBS for 15 min and were washed three times with 1x PBS and permeabilized in 0.5% v/v Triton X-100 in 1x PBS for 10 min, then washed with 1x PBS three times. The tissue slices were then labeled with the MERFISH encoding probes. In brief, the samples were washed with 3 mL of pre-hybridization wash buffer (2x SSC, 30% formamide) for 5 min. The sample was quickly transferred into a hybridization chamber formed on the bottom of a 150-mm petri dish by placing on its bottom a layer of parafilm with a sample-size hole punched in the middle. The parafilm was gently pressed onto the bottom of the petri dish to create a seal. Then, 50 µL hybridization mixture containing 8-10 µM of encoding probes and 1µM of anchoring probes in 2x SSC with 30% v/v formamide, 1 mg/mL yeast tRNA (ThermoFisher Scientific, 15401029), and 10% w/v dextran sulfate (Millipore Sigma, S4030) was added to the petri dish in the middle of the parafilm hole. Then the coverslip was inverted onto the parafilm to completely immerse the tissue section in the droplet of hybridization mixture. The anchor probe was the same as that used previously[1] and had a sequence of /5Acryd/TTG AGT GGA TGG AGT GTA ATT+ TT+ TT+ TT+ TT+ TT+ TT+ TT+ TT+ TT+ T, where /5Acryd/ represents an acrydite modification and T+ indicates locked nucleic acid (ordered from IDT). A wet Kimwipe was placed inside the petri dish to maintain humidity. The petri dish was then closed, sealed with parafilm, and incubated for 36 h at 37 °C in a humidified oven. The coverslip was then removed from the hybridization chamber, placed sample side up, and washed 2x with post-hybridization wash buffer (30% v/v formamide in 2x SSC) for 30 min in a 47 °C oven. Samples were then washed in 2x SSC at room temperature for 5 min and incubated with WGA-oligo conjugate (16 µg/mL in 2x SSC) at room temperature for 1 h, and washed 3x with 2XSSC for 10min at room temperature. Then the coverslip was incubated two times with degassed hydrogel solution consisting of 4% 19:1 acrylamide/bis-acrylamide (Bio-Rad, 1610144) in 50 mM Tris-HCl (ThermoFisher Scientific, 15568-025), 300 mM NaCl (ThermoFisher Scientific, AM9759), 0.03% w/v ammonium persulfate (Millipore Sigma, 215589), and 0.25% v/v tetramethylethylenediamine (TEMED; Sigma, T7024). 50 µL of degassed hydrogel solution was added to cover the tissue section, a 22 mm Gel-Slick (Lonza, 50640) coated coverslip was carefully placed on to the top of the tissue section and incubated for at least 1.5 h to allow polymerization. The 22 mm coverslip was then gently removed with a razor blade. The sample was then incubated in digestion buffer consisting of 1:100 proteinase K (NEW ENGLAND BIOLABS, P8107S) in 0.25% v/v Triton-X (Millipore Sigma, T8787), 2% v/v Sodium Dodecyl Sulfate (SDS; ThermoFisher Scientific, AM9823) in 2x SSC. The sample was incubated at 37 °C in a humidified oven for 18-24 h. The sample was then washed 3x with 2x SSC for 30 min at room temperature. The sample was then either stored in 2x SSC at 4 °C for no more than 3 days or immediately stained with readout probes for imaging.

**Preparation of cardiac tissue sections for histology using Sirius Red/Fast Green collagen staining**

To evaluate collagen deposition, histological staining was performed on cardiac tissue sections located adjacent to those used for MERFISH imaging. Each tissue section was fixed in 4% freshly diluted paraformaldehyde (Electron Microscopy Sciences, 15714), washed in 1x PBS, and stained using the Sirius Red/Fast Green Collagen Staining Kit (Chondrex Inc., Cat. #9046) according to the manufacturer's instructions. Briefly, sections were incubated with a dye solution containing Direct Red 80 (Sirius Red), which selectively stains collagen, and Fast Green FCF, which labels non-collagenous proteins. Following staining, slides were rinsed, dehydrated through graded ethanol, cleared, and mounted. Whole-slide images were acquired using a slide scanner (Olympus Slide Scanner VS120) under brightfield illumination. These images were used for spatial correlation with MERFISH data.

**MERFISH imaging system**

MERFISH imaging was performed on a custom built microscope and fluidics system as described previously.[2] Briefly, the home-built microscope include a Nikon Ti2 microscope body with an high-magnification oil-immersion objective (Nikon, 60x CFI PlanApo lambda) for MERFISH imaging and a low-magnification air objective (Nikon, 10X CFI PlanApo lambda) for sample overview. Sample position was controlled by a motorized XY scanning stage (Marzhauser, SCAN IM 130×85), and auto-focus was controlled by a piezo-based objective nano-positioner (PI PD72Z2CAQ, 250 µm Travel Range). Illumination was provided by a laser light engine (Lumencor, CELESTA quattro VCGRnIR, 90-10713; with 0.8 mm diameter core optical fiber) coupled into the microscope via an epi-illumination module for wide-field illumination (Lumencor, 82-10154). The illumination was reflected to the sample by a pentaband dichroic mirror (Lumencor, 10-10858), and stray illumination was removed from the emitted fluorescence by a pentaband emitter (Lumencor, 10-10857) mounted in the microscope filter cube. The fluorescence emission of the sample was transmitted onto a sCMOS camera (Hamamatsu, ORCA Flash 4.0 V3). The focus of the microscope was maintained using a custom-built autofocus system in which a laser (Thorlabs, LP940-SF30, 940 nm) was split into two beams via polarization, coupled into the microscope via a stage-up kit (Nikon, TI2-LA-SU) containing a filter cube with a dichroic (Thorlabs, DMSP900R) and a long-pass filter (Thorlabs, FEL0900), reflected from the sample-coverslip interface, and imaged on a CMOS camera (Thorlabs, CS165MU). Changes in focus resulted in movements of the imaged beams, providing a signal for closed-loop focus control via the objective nano-positioner. The 60x magnification of this microscope system produced a pixel size of ~107 nm.

The tissue sample was assembled into a commercial flow chamber (FCS2, Bioptechs) with a 0.75-mm-thick flow gasket (Bioptechs DIE F18524; 1907-100), and fluid flow was controlled by a peristaltic pump (Gilson, MP1). The specific buffer drawn across the sample at any given time was controlled by configuring of a set of daisy-chained, computer-controlled 12-port valves (IDEX, EZ1213-820-4). These valves were connected to a custom-built flow manifold and sipper system in which individual stainless-steel needles (Hamilton, 22016-01) were submerged into either 50-mL falcon tubes or different wells of a 24-deep-well plate, into which the various buffers were placed. The microscope and flow system were collectively controlled via custom-built software (github.com/ZhuangLab/storm-control).

**MERFISH imaging**

MERFISH imaging was carried out in 8 sequential imaging rounds in which 2 bits (i.e. readout probes) were imaged in each round. These imaging rounds were separated by fluidics programs in which the sample labeling was reset through cleavage of the disulfide bonds on the hybridized readout probes, followed by hybridization of a set of new readout probes to the sample. To prepare a sample for MERFISH imaging, 2x SSC was first removed. The sample was then stained with 3 mL of readout hybridization/wash buffer which comprised 2x SSC with 10% v/v ethylene carbonate (Alfa Aesar, A15735-36), 0.1% v/v Triton-X (Millipore Sigma, T8787), and 3 nM of two readout probes associated with the first round of imaging and a readout probe for the WGA cell boundary label. The sample was stained with this readout probe mixture for 15 min at room temperature and then incubated with DAPI (1ug/ml in 2x SSC) for 10 min at room temperature. Finally, the sample was washed 3x with 2x SSC and transferred to the microscope for imaging.

To prepare the fluidics for imaging, a readout probe plate was prepared in which individual wells of a 24-deep-well plate (Westnet, 504361) were each filled with 5 mL of the readout hybridization buffer described above, with each well containing a different pair of readout probes associated with a different imaging round. This readout probe plate was loaded into the custom manifold and sipper system described above. Another set of four buffers were prepared and loaded into 50-mL falcon tubes: (1) a cleavage buffer consisting of 45 mM Tris(2-carboxyethyl)phosphine hydrochloride solution (Millipore Sigma, 0.5M pH7.0, 646547-10X1ML) in 2X SSC, to cleave the disulfide bond and extinguish the fluorescence signal after each round of imaging; (2) imaging buffer consisting of 50 µM Trolox-quinone, recombinant protocatechuate 3,4-dioxygenase (rPCO; 1:500, OYC Americas, 46852004), and 5 mM protocatechuic acid (Millipore Sigma, 37580-25G-F) in 2x SSC, pH adjusted to 7.0 with 1M NaOH. Trolox-quinone was made via UV incubation at 254 nm of 2 mM Trolox solution; (3) readout hybridization/wash buffer described above, used to remove stray readout probes after their hybridization; (4) 2x SSC, used to wash away excess cleavage buffer prior to the hybridization of the next set of readout probes.

A fully automated MERFISH measurement began with the selection of the desired fields-of-view (FOV) associated with full coverage of the mouse heart section on the coverslip. These FOVs were selected by creating a low-resolution tiled mosaic of the sample using the 405-nm illumination to image the DAPI-stained sample with the 10x objective. For each of these FOVs, the 60x objective was used to collect a Z-stack comprised of seven 1.5-µm-spaced Z-planes (2048 pixels X 2048 pixels each) in the 735-nm and 635-nm and channels. These two excitation channels selectively excited the Alexa750 and Cy5 fluorophores associated with the the readout probes in the current imaging round. The microscope then collected a single image of the fiducial beads attached to the coverslip (535-nm illumination) to allow registration of images from the same FOV across different imaging rounds. In the first round of imaging, an additional Z-stack was collected for the WGA cell boundary stain using 473-nm illumination, and DAPI-stained nuclei with 405-nm illumination. When all images for all FOVs had been acquired, the flow system was used to immerse the sample in 2 mL of cleavage buffer for 15 min, then 2x with 2 mL of 2x SSC for 3 min, then 2 mL of the next readout hybridization mixture for 15 min, then 2 mL readout hybridization/wash buffer for 10 min, and then 2 mL of imaging buffer for 3 min. The sample was then imaged, as described above, and this process was repeated for a total of 8 imaging rounds. Typical illumination powers at the laser head were 773 mW of 735-nm light, 250 mW of 635-nm light, 30 mW of 535-nm light, 10 mW of 473-nm light, and 20 mW of 405-nm light.

### *3. Data Analysis*

#### Image processing and MERFISH decoding

All MERFISH image analysis was conducted using a custom implementation of the MERlin pipeline (github.com/ZhuangLab/MERlin). Fiducial beads were localized in each field of view (FOV) across imaging rounds to compute affine transformations for correction of drift and chromatic aberration. Preprocessing included high-pass filtering ($\sigma$ = 2.0 pixels) to remove background, followed by Lucy–Richardson deconvolution (10 iterations, Gaussian PSF with $\sigma$ = 1.4 pixels and filter size = 9 pixels) to enhance spot definition. A final low-pass filter ($\sigma$ = 0.6 pixels) was applied to mitigate slight shifts between imaging rounds. A total of 10 iterative optimizations were performed to determine optimal intensity weightings across color channels and imaging rounds using 50 randomly selected FOVs per cycle and a minimum RNA spot area of 5 pixels. Chromatic correction was updated at each step. Barcode decoding was performed by comparing each pixel's weighted intensity vector to expected barcode profiles using a Euclidean metric, with a single-bit error threshold to exclude ambiguous assignments. Barcodes were assigned to pixels and decoded independently for each Z-plane. The decoded data were cropped 89 pixels at every edge of each FOV to remove overlapped regions. Putative RNA molecules were filtered using minimum area of 4 pixels, intensity threshold of 0.9, and a target barcode misidentification rate of 0.5%. Final barcodes, including spatial coordinates (local and global x/y/z), FOV index, and barcode ID, were exported in csv format for downstream spatial and single-cell analysis.

**Cellouette cell segmentation**

Overview: Cellouette consists of three parallel segmentation workflows – Cellouette_s1 (custom WGA-based s1 model), Cellouette_s2 (custom WGA-based s2 model), and Cellpose_DAPI (Cellpose model) – that are subsequently integrated to take advantage of the unique capabilities of each workflow. Of the three workflows, Cellouette_s1 performs best on the large longitudinal CMs; Cellouette_s2 performs best on the transverse CMs; and Cellpose_DAPI performs best when capturing the geometries of the small, non-CM cells in the cardiac tissue section.

Preprocessing and stitching of WGA Images: To generate high-quality mosaic images of WGA-stained cardiac tissue, all FOVs were first corrected for uneven illumination using the flat-field correction function in the sdt-python package[4]. This step compensated for brightness variations across the images. To stitch a seamless whole-tissue mosaic, a gradient-based blending strategy was applied to each FOV, which had 10 μm overlaps with neighboring FOVs (above, below, left, and right). Within these overlapping regions, pixel intensities were blended using a gradient-based weighting function, creating smooth transitions across FOV boundaries and minimizing edge artifacts. Then the whole stitched WGA images were downsampled from 16-bit to 8-bit precision, downscaled (s1 resolution: downscaling by factor of 2; or s2 resolution: downscaling by factor of 4) and converted into multi-resolution Zarr arrays for efficient downstream cell segmentation.

Foreground-Background Model Training and Prediction: To differentiate tissue from background in WGA-stained cardiac tissues, a custom lightweight 2D U-Net model[5] was trained on manually annotated images with binary foreground-background labels. The model was trained using data augmentation and a weighted loss function to address class imbalance. After segmentation, the trained model was used to generate foreground masks, which were then applied in post-processing to remove masks segmented in non-tissue regions. The model was trained for 100,000 iterations using the Adam optimizer (learning rate = $1X10^{-4}$), with batch size = 10.

2D->3D Model Training: To generate instance segmentations of cells in WGA-stained cardiac tissue, a two-step 2D-to-3D pipeline was implemented as previously described[6]. Direct-neighbor affinity graphs (affinities) were used to represent cell boundaries by capturing the connectivity or affinity value of each pixel to its immediate neighbor[7]. This representation, in contrast to binary boundary masks, helped resolve ambiguities in voxel partitioning between instances, particularly when adjacent cells shared boundaries.

Briefly, the first step of the 2D-to-3D pipeline involved training a separate 2D U-Net model to predict dense 2D affinities, with 2D local shape descriptors (LSDs) included as an auxiliary learning task to support instance segmentation. The training dataset consisted of 36 dense and 46 sparse manually annotated images, totaling 11320 labeled cell masks. Affinity labels were generated along 2D spatial axes, and the model was trained using a weighted mean squared error loss with dynamic label balancing and data augmentation.

During training, predicted affinity maps and associated outputs were periodically saved as Zarr files for visual inspection and quality control. These snapshots included raw images, ground truth labels, predicted affinities, and training weights, enabling ongoing monitoring of model performance. The model was trained for 500,000 iterations using the Adam optimizer (learning rate = $1 \times 10^{-4}$) with a batch size of 10.

In the second step of the pipeline, a lightweight 3D U-Net was trained to predict 3D affinities from stacked 2D affinity maps. This network was trained on synthetic 3D labels and received only the 2D affinity predictions generated by the 2D network as input. Once the 2D network produced dense affinity maps across all tissue sections in a volume, the 3D network was applied to generate a full 3D affinity volume, which was used for downstream volumetric segmentation.

*3D Segmentation with Hierarchical Agglomeration:* Predicted 3D affinities were converted into segmented cell instances using a watershed-based hierarchical agglomeration[8]. Initial fragments were generated by performing maxima-seeded watershed on the affinities, in a block-wise manner using Daisy[9]. Low-affinity fragments were filtered out by thresholding based on average affinity values. Edges connecting neighboring fragments were assigned a weight based on the affinity values between the fragments, forming a weighted graph. Edges of the graph were then hierarchically merged using the waterz algorithm (https://github.com/funkey/waterz) up to a merge threshold of 0.10 and a mean affinity-based scoring function. The final segmentation volumes were saved as 3D Zarr arrays.

*Post-processing of 3D Segmentation:* Final cell segmentations were refined through a series of post-processing steps. First, segmentations were masked using the predicted tissue foreground to remove non-tissue regions. Segments that were only segmented on a single Z-slice were excluded. Small holes within segmented cells were filled using morphological binary closing, and boundary pixels between touching cells were removed with a local neighborhood-based filtering step.

*DAPI segmentation (Cellpose_DAPI):* Briefly, stitched DAPI image mosaics were downscaled to s2 resolution and segmented using the Cellpose[10] 2.0 'nuclei' model (diameter = 13), followed by morphological erosion to reduce halo artifacts. These segmentations provided accurate masks for small non-cardiomyocyte (non-CM) cells.

*Integration of segmentation workflows*: To leverage the unique segmentation performance of each workflow, Cellouette_s1, Cellouette_s2, and Cellpose_DAPI were integrated to create a final segmentation with optimal performance across heterozygous cell morphologies. Cellouette_s1 performs best on the large longitudinal CMs; Cellouette_s2 performs best on the transverse CMs; and Cellpose_DAPI performs best when capturing the geometries of the small, non-CM cells in the cardiac tissue section.

The Cellouette_s2 segmentations served as the foundation, with large longitudinal cardiomyocyte masks (>14,000 pixels^2) substituted with corresponding masks from Cellouette_s1, downsampled to match the s2 resolution. These integrated WGA-based segmentation masks were then compared against masks from Cellpose_DAPI. WGA masks were replaced with the corresponding DAPI mask when >50% of the WGA area was overlapped by the DAPI mask. This approach combined the increased accuracy of Cellouette_s1 and Cellouette_s2 for the majority of cells, particularly CMs, together with Cellpose_DAPI providing secondary predictions for smaller non-CM cells with clearer nuclear signal. The optimized and integrated segmentations were saved in 3D Zarr format for downstream analysis.

**Partitioning detected transcripts to segmented cells**

Transcripts (detected barcodes) were assigned to cells using a pixel-based approach. For each transcript, the corresponding pixel coordinate in the segmentation mask was indexed to determine the cell ID at that coordinate (x, y and z). Transcripts falling on background pixels were excluded from cell-based analysis. Gene counts for each cell were aggregated and exported as a cell-by-gene expression matrix. In addition, cell metadata was exported as a separate file that included volume, centroid, bounding box, and z-plane locations of each cell.

**Evaluation of cell segmentation performance**

To quantify the performance of Cellouette relative to other segmentation approaches, 10 FOVs not included in the training or validation sets were manually annotated and used as the ground-truth for testing segmentation performance. The commonly used segmentation algorithms of nuclear watershed, Cellpose[11], and Cellpose SAM[12] were each used to generate cell segmentation predictions from s2 image data.

For the watershed approach, scikit-image was used to generate segmentation masks from DAPI image data. The custom-trained Cellpose 2.0 model was trained on the same manually-annotated training dataset used to train *Cellouette*. Cellpose SAM was used as described[12].

All segmentation predictions were post-processed to remove boundary pixels between touching cells. Three metrics were used to benchmark each segmentation method: *(1) Coverage:* Percentage of transcripts partitioned to cells in ground truth that were partitioned to cells using the given segmentation approach; *(2) Transcripts merged:* Transcripts found in a single cell in the given segmentation approach that were part of multiple cells in the ground truth, given as a percentage of total partitioned transcripts in the ground-truth data. *(3) Transcripts split*: Transcripts found in a single cell in the ground truth data that were part of multiple cells in the given segmentation approach, given as a percentage of total partitioned transcripts in the ground-truth data.

**Mutually exclusive marker expression analysis**

To validate the accuracy of cell segmentation, co-expression of mutually exclusive cell-type marker genes within a single cell was analyzed. Specifically, gene pairs representing four major cardiac cell types were examined: *Ttn* (cardiomyocytes), *Col1a1* (fibroblasts), *Pecam1* (endothelial cells), and *Csf1r* (immune cells). For each possible pair of these genes, transcript counts were volume-normalized at the single-cell level. Cells with zero expression for both genes in a pair were excluded from the analysis. The expression of one gene was plotted against the other and co-expression distributions were visualized using two-dimensional histograms that used a logarithmic color scale to represent the number of cells within each bin. Strong mutual exclusivity between cell-type-specific markers — indicated by minimal overlap in expression within individual cells — served as a quality control metric, confirming segmentation accuracy and low levels of transcript mixing between neighboring cells.

**Single-cell unsupervised clustering**

Volume-normalized and log-transformed gene expression values were processed through the Seurat[13] unsupervised clustering pipeline, to identify major cell populations based on their transcriptomic profiles. Two of the genes in our panel (Wfikkn1, Gdf15) displayed a non-specific (likely artifactual) transcript localization pattern that was not confined to the boundaries of the tissue section and were therefore excluded from downstream analysis. Using the remaining 126 genes, expression data from six mouse hearts (sham and TAC, N = 3 per condition) were integrated using Principal Component Analysis (*RunPCA*), followed by batch correction with Harmony (*RunHarmony*) using a theta value of 2 across the top 100 principal components. The batch-corrected embeddings were used to construct a shared nearest neighbor graph (*FindNeighbors*) based on the top 10 Harmony-adjusted dimensions. Clustering was performed using the Leiden algorithm (*FindClusters*), and results were visualized using Uniform Manifold Approximation and Projection (*RunUMAP*) applied to the same 10 dimensions. UMAP parameters included a minimum distance of 0.01, 60 neighbors, and a spread of 0.5.

**Removal of artifact in the Sham3 dataset**

During quality control, we detected aberrant spatial expression patterns in a subset of genes in the Sham3 sample, likely arising from an imaging defect in one or more MERFISH imaging rounds. To characterize and correct for these artifacts, we binned transcript counts into a fixed spatial grid and quantified gene expression across the tissue section. This approach revealed an artifactual transcript loss for several genes in part of the tissue, with a boundary clearly aligned with the microscope imaging direction. Among affected genes, Pln showed the most pronounced spatial skew and was therefore used to define a linear boundary demarcating the artifact-affected region. This boundary was used to crop the original dataset, and only the retained portion was used for downstream analysis. For the cell population analysis shown in Figure 1F, we imputed the total cell populations in Sham3 by assuming that cell populations were spatially homogeneous across the tissue. We thus scaled the post-crop retained cell populations as: $N_{total} = N_{retained} * S$, where $S = V_{total}/V_{retained}$. $V_{total}$ and $V_{retained}$ are the original and post-crop retained volumes of the tissue, respectively.

**Cell type annotation**

Cell type annotation was carried out through two iterative rounds of clustering and marker-based refinement. In the first round, clustering at resolution0.5 identified 10 initial clusters. One cluster was flagged as a likely population of doublets based on two independent observations: (1) Intermediate UMAP positioning: this cluster appeared as a bridging subpopulation between two well-separated clusters; (2) upon visual inspection, many of these cells displayed irregular segmentation including segmented interstitial spaces that lacked distinct cellular boundaries, indicative of segmentation artifacts. This putative doublet population was removed, and a second round of clustering was performed at a lower resolution (0.3), yielding 8 clusters. Subsequently, the expression of canonical marker genes was used to merge clusters into 4 major cell types: Cardiomyocytes (CM), Endothelial Cells (EC), Fibroblasts (FB), and Immune cells (IC).

**Cell shape analysis and cardiomyocyte morphotype classification**

To characterize cell morphology, 2D geometric features were extracted from a representative Z-slice of the segmentation mask using the function *skimage.measure.regionprops_table* from *scikit-image*[14]. For each segmented cell, we extracted centroid coordinates, major and minor axis lengths, eccentricity, solidity, perimeter, orientation, area, and extent. The aspect ratio (minor axis length /major axis length) and form factor (4π area /perimeter²) were calculated to assess cell shape (circular or elongated). Due to the large size and rod-like geometry of cardiomyocytes, segmented cardiomyocytes could have distinct shapes depending on their orientation within the tissue. To avoid introducing bias in downstream analyses of cardiomyocyte size and morphological changes, orientation-based classification was applied to separate longitudinal and cross-sectional cardiomyocytes: Cross-sectional cardiomyocytes were defined as cells with aspect ratio ≥ 0.6 and form factor ≥ 0.6, indicating compact, rounded cells consistent with transverse sections of cardiomyocytes. Longitudinal cardiomyocytes were identified as cells with aspect ratio < 0.2, eccentricity > 0.98, and an area > 500 μm², indicating elongated cells consistent with longitudinal sections of cardiomyocytes.

**Estimation of cardiomyocyte orientation angle**

To generate the cardiomyocyte orientation graph in Figure 1I, we derived 3D CM orientation vectors through a multistage process. For each CM, we first used the 2D CM cell segmentation masks to determine for the CM's 2D centroid horizontal displacements (dx,dy) across the z planes within the tissue section (dz = ~1.5μm between imaging planes). We then computed the magnitude of each CM's average dx and dy per z plane, yielding the CM's average centroid displacement across z planes. We then identified the CM's largest 2D segmentation mask (by area) across the z planes in which the cell appeared, and measured the lengths of that mask's major and minor axes. We converted these axis lengths to an eccentricity value for every CM. We then fit a LOESS curve to define cell centroid displacement as a function of cell eccentricity, and computed the signed difference between actual centroid displacement and LOESS-predicted centroid displacement for cells. We then converted this difference to a probability for in-plane orientation of that cell, by normalizing differences to a [0,1] range, and reversed the scale so that higher normalized values correspond to higher probability of the cell being oriented within the plane. We then smoothed this probability by applying a sigmoid transformation. Finally, we reversed the probability scale once again and scaled each CM's dz by its smoothed, reversed in-plane probability.

**Cardiomyocyte subclustering**

To characterize cardiomyocyte (CM) heterogeneity, the CM population was isolated and subjected to iterative subclustering. To avoid inclusion of cells with insufficient transcript counts for accurate analysis, cells with a total volume ≤ 500 $\mu m^3$ were excluded, thus removing small, fragmented regions that typically corresponded to cardiomyocyte tips. Additionally, four genes with a high degree of subcellular variation in density — *Xirp2*, *Ccn2*, *Slc16a1*, and *Slc2a4* — were excluded to minimize artifacts caused by transcript enrichment at specific subcellular locations relative to the sectioning plane. This filtering ensured that clustering was driven by transcripts more evenly distributed throughout the cytoplasm. CMs were then clustered using the Leiden algorithm with a resolution of 0.25, resulting in five initial subclusters. To refine these into biologically meaningful groups, pairwise differential expression analysis was performed using DESeq2[15]. Genes were considered significantly differentially expressed if they met the following criteria: absolute $\log_2$ fold change > 1.5, adjusted p-value < 0.05, and expression detected in more than 50% of cells in at least one of the compared clusters. Based on differential expression of *Acta1* and collagen genes (*Col1a1*, *Col1a2*, *Col3a1*), the five subclusters were consolidated into three transcriptionally distinct CM subtypes: Subcluster 1 (CM1): *Acta1*$^{-}$*/ Col1a1/1a2/3a1*$^{-}$, Subcluster 2 (CM2): *Acta1*$^{+}$*/ Col1a1/1a2/3a1*$^{-}$, Subcluster 3 (CM3): *Acta1*$^{+}$*/ Col1a1/1a2/3a1*$^{+}$. These three CM subtypes were used for downstream analyses and biological interpretation.

**SpatialScrub automated doublet removal**

A custom workflow that we named SpatialScrub was used to remove (i.e. scrub) suspected doublet cells based on their spatial distribution of transcripts. The scrubbed lists of cells were used for all the downstream analyses presented in Figures 2-5 and accompanying Supplemental Figures.

Upon visual inspection we found that there was a small number (<2%) of CMs that had an unusually high subcellular density of collagen transcripts that were spatially clustered within a small part of the cell. These subcellular regions lacked canonical CM markers. We therefore identified these CMs as CM-FB doublets. To remove these CM-FB doublets, we used SpatialScrub. In brief, we used the subcellular spatial coordinates of collagen transcripts within the cardiomyocyte cell geometry. We identified clusters of collagen transcripts by using the DBSCAN algorithm. Specifically, a CM-FB doublet was defined as a CM containing a cluster of collagen transcripts as detected using the scikit-learn DBSCAN function with parameters eps=2 μm (maximum distance between transcripts), and min_samples=10 (minimum of 10 transcripts). CMs with at least one subcellular cluster of collagen transcripts were removed from all downstream analyses presented in Figures 2-5 and accompanying Supplemental Figures.

Additionally, we identified a small number of immune cells that were split into multiple segmentation masks, meaning one cell was defined by multiple nearby but non-overlapping segments (geometries). Inspection of the transcript distribution showed that one geometry typically contained IC marker genes, while another contained FB markers. Furthermore, examination of the 3D cell masks revealed that, often in a single Z-plane, two distinct cells had been incorrectly segmented as one. These cells were removed from all downstream analyses presented in Figures 3-5 and accompanying Supplemental Figures.

**Spatial patterns of cardiomyocyte-specific gene expression**

<u>Spatially variable genes analysis:</u> To enable spatially variable gene analysis within the cardiomyocytes (CM), the cell-by-gene expression was filtered to include only genes with log-transformed, mean volume-normalized expression in CM greater than 'blank' barcodes (>0.35) in TAC1. For genes that remained, volume-normalized CM gene expression was mean-rasterized into a hexagonal grid using SEraster[16]. Briefly, given the cell-by-gene expression matrix and the cell centroid positions of CM in the TAC hearts, the coordinate spaces of the datasets were binned into evenly spaced hexagons with 171μm resolution and for each hexbin, the mean volume normalized expression was calculated from the CM with centroids within the hexbin boundaries (*rasterizeGeneExpression*).

The MERINGUE[17] package's implementation of Moran's *I* was used to identify spatially variable genes. Moran's *I* was calculated for each gene's log-transformed, mean-rasterized volume-normalized expression with pseudocount +1 (*getSpatialPatterns*). For creating the binary adjacency weight matrix, a filtering distance of 172μm was used to encode the neighbor-relationships for the rasterized hexbins (*getSpatialNeighbors*). Genes were identified as spatially variable if its observed Moran's *I* had an adjusted p-value < 0.05 and if the gene had a pattern driven by more than 5% of the hexbins, which was calculated as the percentage of hexbins that had statistically significant (p-value < 0.05) local indicator of spatial association scores (*filterSpatialPatterns*).

<u>*Pattern identification via grouping spatially correlated genes:*</u> To group the spatially variable genes with similar patterns of spatial heterogeneity into spatial patterns groups, a spatial cross-correlation matrix was constructed by calculating the Pearson correlation of the hexbin volume-normalized expression between all pairwise combinations of spatially variable genes. The resulting matrix was analyzed through MERINGUE using Ward's hierarchical clustering and dynamic tree cutting with minimum cluster size of 0 and *deepSplit* parameter set to 0 (*groupSigSpatialPatterns*) to group genes into spatial pattern groups.

<u>*Reproducibility of patterns across TAC hearts:*</u> To determine if the observed gene expression spatial patterns A and B were reproducible across the TAC hearts, only the genes within either pattern A or B in the first TAC heart (TAC1) were used to construct spatial cross-correlation matrices of hexbin volume-normalized expression for the second and third TAC hearts (TAC2 and TAC3). The resulting matrices were also analyzed via hierarchical clustering and dynamic tree cutting to group genes into spatial pattern groups.

For the genes within a pattern group, a mean expression z-score was calculated by performing z-score normalization on the hexbin volume-normalized expression of each gene within the pattern and averaging across the genes. Jaccard similarity was used to measure the similarity of pattern groups across the TAC hearts with regards to which genes are members of each group. For a pair of spatial patterns, the Jaccard similarity was measured as the number of genes in the intersection of the groups divided by the number of genes in the union of the groups.

<u>*Correlation of gene expression patterns with CM states and fibroblast abundance:*</u> To relate the spatial gene expression patterns to the CM states, the CM subcluster classifications were sum-rasterized into a hexagonal grid using SEraster[16]. Briefly, given the CM subcluster labels (CM1, CM2, CM3) and the cell centroid positions for CM in the TAC hearts, the coordinate spaces of the datasets were binned into evenly spaced hexagons with 171μm resolution and for each hexbin, the total number of CM in each subcluster was calculated from the CM with centroids within the hexbin boundaries (*rasterizeCellType*). The proportion of each CM subcluster in each hexbin was then calculated as the number of each CM subcluster divided by the total number of CM within the hexbin. Pearson's correlation was calculated between the proportion of each CM subcluster and the expression z-score of each pattern group. The fibroblasts were also rasterized into hexbins and the pearson's correlation was calculated between the number of fibroblasts within each hexbin and the expression z-score of each pattern group.

**Cell neighborhood analysis**

<u>*Cell neighborhood graph construction:*</u> To characterize spatial cellular interactions between neighboring cells in cardiac tissue, network graphs were constructed using NetworkX (https://networkx.org/) Each graph node represented the centroid of a segmented cell from a representative 2D Z-plane and was annotated with node attributes including cell ID, cell type, subcluster identity, etc. Cell boundary coordinates were extracted from segmentation masks and stored as Shapely polygon objects (https://shapely.readthedocs.io/). For every pair of cells, the minimum Euclidean distance between their polygon boundaries was computed. If this distance was less than a specified threshold, the cells were considered neighbors, and an edge was added between their corresponding nodes.

<u>*Distance threshold optimization:*</u> To determine an optimal threshold for defining cell neighborhoods, a series of network graphs were generated using distance thresholds ranging from 1.5 μm to 4.5 μm, in 0.25 μm increments. For each threshold, a variability score was computed by comparing its graph to its adjacent threshold graphs (±0.25 μm). Specifically, for each threshold-adjacent graph, for every node the absolute difference in its number

of neighbors between the threshold and adjacent graph was calculated. These absolute differences were summed across all nodes to give a sum of absolute difference for each adjacent graph. The sums of absolute difference from both adjacent graphs were averaged to define the variability score for that threshold. This analysis was conducted independently for both sham and TAC heart datasets. In both cases, the 3 µm threshold produced the lowest variability score, indicating it was the most stable and biologically representative distance for defining cell-cell proximity. The final network graphs were thus constructed using a 3 µm distance threshold and were used in subsequent analyses.

*Node-edge quantification:* The finalized network graphs were used to quantify cell-type-specific spatial connectivity. For every cardiomyocyte node, the number of its edges connected to its neighbor nodes of each major cell type (cardiomyocytes, fibroblasts, endothelial cells, immune cells) were totaled to give the total number of cardiomyocyte-specific interactions across the entire network graph. The total number of interactions with each cell type were normalized by the total number of cardiomyocyte nodes to compute the average number of cell-type-specific interactions per cardiomyocyte. To further explore spatial organization within cardiomyocyte subpopulations, interactions between each cardiomyocyte subtype with each major cell types were quantified, revealing connectivity features linked to transcriptional states.

**Microenvironment analysis through hexbin construction, quantification, and feature correlation**

*Hexbin grid construction:* To enable spatially resolved analysis of local tissue features and transcriptional relationships with single-cell resolution, cardiac tissue sections were computationally binned into evenly spaced hexagons (short diagonal length of 171 µm) resulting in a hexagonal grid covering the entire cardiac tissue. Each hexbin was assigned a unique ID and referred to by its centroid coordinates. Within each hexbin, various features were quantified including cellular composition by major cell types and subclusters, cell-type-specific gene expression, and fibrosis coverage.

*Hexbin quantification – cellular composition:* The hexbin grid was overlaid on the cell segmentation mask to spatially quantify cellular composition. For each hexbin, the number of overlapping pixels with the cell masks were computed and grouped by cell type to quantify cell-type-specific pixel coverage for the hexbin. Cardiomyocyte (CM) subcluster composition was also quantified per hexbin and later used for CM state-specific enrichment analysis.

*Hexbin quantification – fibrosis coverage:* To quantify local fibrosis within each hexbin, fibrosis staining images from adjacent tissue sections were aligned to corresponding cell segmentation mask data, enabling spatial integration of fibrotic features with MERFISH data. First, histology and segmentation image registration were performed using STalign[19]. Briefly, high-resolution histological image and segmentation mask data were downsampled and then normalized to an intensity range of [0, 1]. The histology images were padded to match the dimensions of the segmentation mask images. Using the STalign graphical interface, shared anatomical landmarks were manually annotated across both sets of images. An initial affine transformation was computed from these landmark points and refined via Large Deformation Diffeomorphic Metric Mapping (LDDMM), treating the segmentation mask as the target and the histological image as the source. This produced a non-linear transformation aligning tissue morphology across modalities, allowing spatial correspondence between segmented cells and fibrotic staining. Second, to identify fibrotic regions, histological images were converted from RGB to HSV color space. Fibrotic pixels were identified by thresholding the HSV image based on lower-bound HSV thresholds (sham sections: [0.87, 0.25, 0.8] , TAC sections: [0.87, 0.1, 0.8]), which captured the fibrotic pixels in collagen-rich regions. Pixels exceeding all three HSV channel thresholds were classified as fibrotic, and a binary fibrotic mask was created with fibrotic pixels set to 1. Third, the hexbin grid was overlaid on the binary fibrotic mask. For each hexbin, the fibrosis coverage was calculated as [no. of fibrotic pixels] / ([no. of pixels containing cells] + [no. of fibrotic pixels]).

*Hexbin quantification – cell-type specific gene expression:* To quantify gene expression for each cell type within each hexbin, transcript coordinates from every cell were mapped onto the hexbin grid. For each hexbin, the transcripts partitioned to each cell were grouped according to the cell's major cell type and CM subcluster identity, enabling quantification of cell-type-specific and CM subcluster-specific gene expression.

For each hexbin, for every gene, the following gene expression features were quantified: total transcript counts per gene (all cells); cell-type-specific transcript counts per gene (CM, EC, FB, IC); CM subcluster-specific transcript counts per gene (CM1, CM2, CM3); gene expression relative to hexbin cell coverage; all-cell density (transcripts per gene normalized by total cell-covered area (counts/µm²)); cell-type-specific density (transcripts per gene normalized by the area covered by each specific cell type or subcluster (counts/µm²)).

*Hexbin feature correlation analysis:* Hexbin-based correlation analysis was performed on one representative dataset from each condition (sham and TAC) to identify spatial relationships between cellular, transcriptional and morphological features. For each dataset, Spearman correlation coefficients and adjusted p-values were calculated for all unique pairs of features across all hexbins, and feature pairs with adjusted p-values < 0.05 were considered statistically significant.

**Cardiomyocyte progression scores**

CM progression scores were calculated from single-cell transcript counts of the FA metabolism genes Cd36, Acsl1, Acadm, and Lpl. For each CM cell with index $i$ and gene $g$, raw transcript counts $c_{i,g}$ were normalized by cellular volume and clipped to the 2$^{nd}$ and 98$^{th}$ percentiles within a tissue section. Volume-normalized and clipped expression values $\tilde{c}_{i,g}$ were then scaled to the unit interval,

$$y_{i,g} = \frac{\tilde{c}_{i,g} - min_j(\tilde{c}_{j,g})}{max_j(\tilde{c}_{j,g}) - min_j(\tilde{c}_{j,g})} \tag{1}$$

and used to compute a per-cell aggregate score by summing across the FA metabolism genes $\mathcal{G}_{FA} = \{\mathrm{Cd36, Acsl1, Acadm, Lpl}\}$,

$$\mathrm{s_i} = \sum_{g \in \mathcal{G}_{FA}} y_{i,g} \tag{2}$$

CM metabolic Progression Scores ($PS_i$) were then calculated by rescaling to the unit interval and inverting the scale so that a lower expression of FA metabolism genes corresponded to a higher progression score:

$$PS_i = 1 - \frac{s_i - min_j(s_j)}{max_j(s_j) - min_j(s_j)} \tag{3}$$

**Cardiomyocyte progression score inference using χNet**

*χNet model overview:* The $\chi$Net model was implemented as a simple fully connected multilayer perceptron (MLP) that predicts for each CM with index $i$, a scalar metabolic score, $\widehat{PS}_i$, based on inputs consisting of cell type-specific expression profiles of local non-CM neighbors.

Local environment parameterization: For each sample, we captured cellular microenvironments using our experimentally determined spatial cell connectivity graph. Cells were represented as nodes in this undirected graph. For a graph $G = (V, E)$ with nodes $v_i \in V$ and a set of edges $E$. A node's cell type label is given by:

$$F_{ct}: v_i \rightarrow \{\mathrm{CM, IC, FB, EC}\} \tag{4}$$

and its volume-normalized gene expression vector is given by

$$F_{exp}: v_i \rightarrow \mathbb{R}^P \tag{5}$$

where $P$ is the number of measured genes.

Let the set of CM cells $V^{CM} = \{v_i \in V | F_{ct}(v_i) = CM\}$. For a single CM cell $cm_i \in V^{CM}$, its neighbor cells at a distance of exactly $d$ hops is:

$$\mathcal{N}_d(cm_i) = \{v_j \in V \mid dist(cm_i, v_j) = d\} \quad (6)$$

where $dist(cm_i, v_j)$ is the shortest-path distance in hops between cells $cm_i$ and $v_j$ in the graph $G$.

Of these neighbor cells, we independently considered each non-CM cell type: immune cell (IC), fibroblast (FB), and endothelial cell (EC). For each non-CM cell-type $NCM \in \{\text{IC}, \text{FB}, \text{EC}\}$, let $V^{NCM} = \{v_j \in V | F_{ct}(v_j) = NCM\}$. The set of non-CM neighbor cells of cell-type $NCM$ at distance $d$ is then:

$$\mathcal{N}_d^{NCM}(cm_i) = \mathcal{N}_d(cm_i) \cap V^{NCM} \quad (7)$$

We formed the expression profile of the non-CM microenvironment surrounding $cm_i$, within a maximum hop distance $D$. We did this by calculating inverse-hop–weighted mean expression vectors $\boldsymbol{\mu}^{NCM}: (cm_i, D) \to \mathbb{R}^P$ summarizing the neighborhood expression profile for each $NCM$ cell type:

$$\boldsymbol{\mu}^{NCM}(cm_i, D) = \frac{\sum_{d=1}^{D} w_d \left(\sum_{v_j \in \mathcal{N}_d^{NCM}(cm_i)} F_{exp}(v_j)\right)}{\sum_{d=1}^{D} w_d \left|\mathcal{N}_d^{NCM}(cm_i)\right|}, \quad w_d = \frac{1}{1+d^2} \quad (8)$$

If no neighbors of type $NCM$ exist within $D$ hops, we set $\boldsymbol{\mu}^{NCM}(cm_i, D) = 0$.

<u>Construction of $\chi$Net inputs:</u> For each dataset, we split CM cell indices $i$ into training and validation partitions, $\mathcal{D}_{\text{train}}$ and $\mathcal{D}_{\text{val}}$, with an 80/20 split. We used the training partition to form a training matrix $\mathbf{X}^{NCM}$ for each non-CM cell-type $NCM$, containing as rows $\boldsymbol{\mu}^{NCM}(cm_{i \in \mathcal{D}_{\text{train}}}, D)$:

$$\mathbf{X}^{NCM} \in \mathbb{R}^{|\mathcal{D}_{\text{train}}| \times P} \quad (9)$$

and fit a PCA model with $K = 10$ components, yielding mean vector $\bar{\boldsymbol{\mu}}^{NCM}$ and loading matrix $\mathbf{U}^{NCM} \in \mathbb{R}^{P \times K}$. The PCA model was fit on the training partition and then applied to training and validation partitions.

Each CM cell $cm_i$'s local cell-type $NCM$ expression vector PCA embedding is defined as:

$$\boldsymbol{v}^{NCM}(cm_i, D) = (\boldsymbol{\mu}^{NCM}(cm_i, D) - \bar{\boldsymbol{\mu}}^{NCM})\mathbf{U}^{NCM} \quad (10)$$

We used these gene expression embeddings to construct the inputs to $\chi$Net, as specified below.

Two different input configurations were used, depending on whether we were predicting based on environmental data from all non-CM cell-types or from a single non-CM cell-type:

1. **All-cell-type model:** the input $\boldsymbol{x}_i$ is a concatenation of three PCA-reduced neighbor expression vectors

$$\boldsymbol{x}_i = [\boldsymbol{v}^{IC}(cm_i, D), \boldsymbol{v}^{FB}(cm_i, D), \boldsymbol{v}^{EC}(cm_i, D)] \in \mathbb{R}^{1 \times 30} \quad (11)$$

2. **Single-cell-type model:** the input $\boldsymbol{x}_i$ is a single PCA-reduced neighbor expression vector

$$\boldsymbol{x}_i = \boldsymbol{v}^{NCM}(cm_i, D) \in \mathbb{R}^{1 \times 10} \quad (12)$$

<u>$\chi$Net model architecture:</u> Given an input vector $\boldsymbol{x}_i$ for CM cell $cm_i$, the $\chi$Net model computes $f_\chi(\boldsymbol{x}_i)$, which consists of a sequence of hidden-layer transformations followed by a linear output representing $\widehat{PS}_i$. We used a fixed MLP architecture with the following layer dimensions:

$$d_{in} \to 50 \to 50 \to 25 \to 10 \to 1$$

where $d_{in}$ is the input dimension (either 30 or 10 depending on input configuration).

Let $\boldsymbol{h}_i^{[\ell]}$ denote the representation of cell $cm_i$ at layer $\ell$, while $\mathbf{W}^{[\ell]}$ and $\boldsymbol{b}^{[\ell]}$ denote the trainable model weights and biases, respectively. We set $\boldsymbol{h}_i^{[0]} = \boldsymbol{x}_i$. Then, for $\ell \in [0,3]$:

$$\boldsymbol{h}_i^{[\ell+1]} = \phi\left(\boldsymbol{h}_i^{[\ell]}\mathbf{W}^{[\ell]} + \boldsymbol{b}^{[\ell]}\right) \tag{13}$$

and the final prediction is

$$\widehat{PS}_i = \boldsymbol{h}_i^{[5]} = \boldsymbol{h}_i^{[4]}\mathbf{W}^{[4]} + \boldsymbol{b}^{[4]} \tag{14}$$

The activation function $\phi(z)$ is the leaky-ReLU:

$$\phi(z) = \begin{cases} z, & for\ z \geq 0 \\ 0.01z, & for\ z < 0 \end{cases} \tag{15}$$

Model training: Given microenvironmental parameterizations of CM cells, $\boldsymbol{x}_i$, and corresponding ground-truth metabolic progression scores, $PS_i$, the $\chi$Net parameters $\theta = \{\mathbf{W}, \boldsymbol{b}\}$ were optimized using Mean Squared Error (MSE) loss:

$$\mathcal{L}_{\text{train}}(\theta) = \frac{1}{|\mathcal{D}_{\text{train}}|}\sum_{i\in\mathcal{D}_{\text{train}}} (\widehat{PS}_i - PS_i)^2 \tag{16}$$

Mini-batch optimization was performed by computing gradients via backpropagation and updating parameters using the Adam optimizer (learning rate $\eta = 10^{-4}$), with a batch size of 50 over 100 epochs. Validation performance was monitored using MSE loss:

$$\mathcal{L}_{\text{val}}(\theta) = \frac{1}{|\mathcal{D}_{\text{val}}|}\sum_{i\in\mathcal{D}_{\text{val}}} (\widehat{PS}_i - PS_i)^2 \tag{17}$$

A fixed seed (42) was used for all PyTorch, NumPy, CUDA, and Python RNGs to ensure reproducibility.

Cell-type and gene importance analysis: We used Shapley values to quantify the contribution of each environmental feature toward model predictions in the validation set $\mathcal{D}_{\text{val}}$ post $\chi$Net training. Two granularities were evaluated:

1. **Cell-type Shapley (used for Figure 5E):** features were the cell-type PCA blocks:

$$\{\boldsymbol{v}^{IC}(cm_i, D), \boldsymbol{v}^{FB}(cm_i, D), \boldsymbol{v}^{EC}(cm_i, D)\}$$

2. **PC-level Shapley (used for Figure 5F):** for a chosen cell type $NCM$, features were the PCA values:

$$\{\boldsymbol{v}^{NCM}(cm_i, D)^{(1)}, \ldots, \boldsymbol{v}^{NCM}(cm_i, D)^{(10)}\}$$

Let $\mathcal{F}$ denote either the set of all features (cell-type or PC-level, depending on whether the all-cell-type or single-cell-type input configuration was used). For a subset $S \subseteq \mathcal{F}$, features in $S$ were left intact while features in $\mathcal{F}\backslash S$ were randomly permuted across $\boldsymbol{x}_i$, for $i \in \mathcal{D}_{\text{val}}$. Let $\boldsymbol{x}_i^{(S)}$ be the resulting perturbed microenvironmental features for CM cell $i$, and let the output of a pre-trained $\chi$Net model, $f_\chi\left(\boldsymbol{x}_i^{(S)}\right)$, be the prediction given this perturbed input $\boldsymbol{x}_i^{(S)}$. The loss for subset $S$ was defined as

$$L(S) = \frac{1}{|\mathcal{D}_{\text{val}}|}\sum_{i\in\mathcal{D}_{\text{val}}} \left(f_\chi\left(\boldsymbol{x}_i^{(S)}\right) - PS_i\right)^2 \tag{18}$$

For each subset $S$ this process was repeated with 10 permutations, ensuring stability of observed loss. The final loss we associated with subset $S$ was the mean loss across the 10 permutations, denoted as $L'(S)$.

The Shapley value for environmental feature $j$ was then

$$\varphi_j = \sum_{S \subseteq \mathcal{F}\{j\}} \frac{|S|!\,(|\mathcal{F}| - |S| - 1)!}{|\mathcal{F}|!} \big(L'(S) - L'(S \cup j)\big) \tag{19}$$

Higher $\varphi_j$ indicates that preserving environmental feature $j$ increased accuracy of predicted CM progression score $\widehat{PS}_i$. Thus, Shapley values identified which neighbor cell-types or individual genes' expression within neighbor cell-types carry the most information regarding CM progression score.

Mapping PC-level Shapley values to gene-level importance: Let $\mathbf{U}^{NCM} \in \mathbb{R}^{P \times K}$ denote the PCA loading matrix for cell type $NCM$. The columns of $\mathbf{U}^{NCM}$ are then $\boldsymbol{u}_1, \dots, \boldsymbol{u}_K \in \mathbb{R}^P$, which contain the gene loadings for each principal component. Let $\varphi_1, \dots, \varphi_K$ be the Shapley importance values obtained from the PC-level analysis.

The gene-level importance vector $\boldsymbol{g}^{NCM} \in \mathbb{R}^P$ is computed as the Shapley-weighted sum of PC loadings:

$$\boldsymbol{g}^{NCM} = \sum_{k=1}^{K} \varphi_k \, \boldsymbol{u}_k \tag{20}$$

In Figure 5F, for each $NCM$ (EC, FB, IC), we display the genes with the top 10 highest values in the corresponding gene-level importance vectors $\boldsymbol{g}^{EC}, \boldsymbol{g}^{FB}, \boldsymbol{g}^{IC}$.

**Visualization with BellaVista**

A customized version of BellaVista[18], built on the napari GUI, was used for interactive visualization of MERFISH data and figure generation. The software included the following features: *(1)* Cardiac tissue visualization with stitched WGA mosaic images in OME-Zarr format optimized for multiscale image browsing. *(2)* Spatial mapping of transcripts: transcripts were visualized using napari point layers, where each gene was assigned a unique color. Transcript positions were derived from globally registered x,y coordinates. *(3)* Spatial mapping of cells: segmented cells were visualized as either napari tracks (cell boundaries) or label layers (cell masks). Each track or label represented an individual cell and was color-coded according to cell type. *(4)* Spatial mapping of cellular connection graph: cell centroids were displayed as napari point layers (nodes) and colored by cell type. Edges were rendered using napari vector layers to connect two neighboring cells if the minimum distance between their boundaries was less than 3 μm.

The BellaVista software and installation instructions can be found on GitHub: BellaVista GitHub Repo. Note that installation of uv is required to run BellaVista. After installing uv (and restarting the terminal), you can run the following commands in a terminal to download and view the Sham1 and TAC1 datasets, respectively:

```
uvx -p 3.12 bellavista --dataset-url "https://bit.ly/KoLabSham"

uvx -p 3.12 bellavista --dataset-url "https://bit.ly/KoLabTAC"
```

### References (Methods)


1. Moffitt, J.R., Hao, J., Bambah-Mukku, D., Lu, T., Dulac, C., and Zhuang, X. (2016). High-performance multiplexed fluorescence in situ hybridization in culture and tissue with matrix imprinting and clearing. Proc Natl Acad Sci U S A *113*, 14456-14461. 10.1073/pnas.1617699113.

2. Moffitt, J.R., Bambah-Mukku, D., Eichhorn, S.W., Vaughn, E., Shekhar, K., Perez, J.D., Rubinstein, N.D., Hao, J., Regev, A., Dulac, C., and Zhuang, X. (2018). Molecular, spatial, and functional single-cell profiling of the hypothalamic preoptic region. Science *362*. 10.1126/science.aau5324.

3. Moffitt, J.R., Hao, J., Wang, G., Chen, K.H., Babcock, H.P., and Zhuang, X. (2016). High-throughput single-cell gene-expression profiling with multiplexed error-robust fluorescence in situ hybridization. Proceedings of the National Academy of Sciences of the United States of America *113*, 11046-11051. 10.1073/pnas.1612826113.

4. Schrangl, L. (2020). sdt-python: Python library for fluorescence microscopy data analysis. Zenodo. https://doi.org/10.5281/zenodo.4604495.

5. Ronneberger, O., Fischer, P., and Brox, T. (2015). U-Net: Convolutional Networks for Biomedical Image Segmentation. held in Cham, (Springer International Publishing), pp. 234-241.

6. Thiyagarajan, V.V., Sheridan, A., Harris, K.M., and Manor, U. (2024). Sparse Annotation is Sufficient for Bootstrapping Dense Segmentation. bioRxiv, 2024.2006.2014.599135. 10.1101/2024.06.14.599135.

7. Turaga, S.C., Briggman, K.L., Helmstaedter, M., Denk, W., and Seung, H.S. (2009). Maximin affinity learning of image segmentation. arXiv:0911.5372. 10.48550/arXiv.0911.5372.

8. Funke, J., Tschopp, F., Grisaitis, W., Sheridan, A., Singh, C., Saalfeld, S., and Turaga, S.C. (2019). Large Scale Image Segmentation with Structured Loss Based Deep Learning for Connectome Reconstruction. IEEE Trans Pattern Anal Mach Intell *41*, 1669-1680. 10.1109/TPAMI.2018.2835450.

9. Nguyen, T., Malin-Mayor, C., Patton, W., and Funke, J. (2022). Daisy: block-wise task dependencies.

10. Stringer, C., and Pachitariu, M. (2022). Cellpose 2.0: how to train your own model. bioRxiv, 2022.2004.2001.486764. 10.1101/2022.04.01.486764.

11. Stringer, C., Wang, T., Michaelos, M., and Pachitariu, M. (2021). Cellpose: a generalist algorithm for cellular segmentation. Nature Methods *18*, 100-106. 10.1038/s41592-020-01018-x.

12. Pachitariu, M., Rariden, M., and Stringer, C. (2025). Cellpose-SAM: superhuman generalization for cellular segmentation. bioRxiv, 2025.2004.2028.651001. 10.1101/2025.04.28.651001.

13. Hao, Y., Stuart, T., Kowalski, M.H., Choudhary, S., Hoffman, P., Hartman, A., Srivastava, A., Molla, G., Madad, S., Fernandez-Granda, C., and Satija, R. (2024). Dictionary learning for integrative, multimodal and scalable single-cell analysis. Nat Biotechnol *42*, 293-304. 10.1038/s41587-023-01767-y.

14. van der Walt, S., Schonberger, J.L., Nunez-Iglesias, J., Boulogne, F., Warner, J.D., Yager, N., Gouillart, E., Yu, T., and scikit-image, c. (2014). scikit-image: image processing in Python. PeerJ *2*, e453. 10.7717/peerj.453.

15. Love, M.I., Huber, W., and Anders, S. (2014). Moderated estimation of fold change and dispersion for RNA-seq data with DESeq2. Genome Biol *15*, 550. 10.1186/s13059-014-0550-8.

16. Aihara, G., Clifton, K., Chen, M., Li, Z., Atta, L., Miller, B.F., Satija, R., Hickey, J.W., and Fan, J. (2024). SEraster: a rasterization preprocessing framework for scalable spatial omics data analysis. Bioinformatics *40*. 10.1093/bioinformatics/btae412.

17. Miller, B.F., Bambah-Mukku, D., Dulac, C., Zhuang, X., and Fan, J. (2021). Characterizing spatial gene expression heterogeneity in spatially resolved single-cell transcriptomic data with nonuniform cellular densities. Genome Res *31*, 1843-1855. 10.1101/gr.271288.120.

18. Coles, A.M., Liu, Y., and Kosuri, P. (2025). BellaVista: Open-Source Visualization for Imaging-Based Spatial Transcriptomics. bioRxiv, 2025.2001.2007.631783. 10.1101/2025.01.07.631783.

19. Clifton, K., Anant, M., Aihara, G., Atta, L., Aimiuwu, O.K., Kebschull, J.M., Miller, M.I., Tward, D., and Fan, J. (2023). STalign: Alignment of spatial transcriptomics data using diffeomorphic metric mapping. Nat Commun *14*, 8123. 10.1038/s41467-023-43915-7.
